# NMR study of the interaction between MinC and FtsZ and modeling of the FtsZ:MinC complex

**DOI:** 10.1101/2024.03.31.587266

**Authors:** Luciana E. S. F. Machado, Patricia Castellen, Valdir Blasios, Helder Veras Ribeiro Filho, Alexandre W. Bisson-Filho, Jhonatan S. Benites-Pariente, Maria Luiza Caldas Nogueira, Mauricio Sforça, Rodrigo Vargas Honorato, Paulo Sérgio Lopes de Oliveira, Roberto K. Salinas, Ana Carolina de Mattos Zeri, Frederico J. Gueiros-Filho

## Abstract

The Min system is a key spatial regulator of cell division in rod-shaped bacteria and the first FtsZ negative modulator to be recognized. Nevertheless, despite extensive genetic and in vitro studies, the molecular mechanism used by MinC to inhibit Z-ring formation remains incompletely understood. The crystallization of FtsZ in complex with other negative regulators such as SulA and MciZ has provided important structural information to corroborate in vitro experiments and establish the mechanism of Z-ring antagonism by these modulators. However, MinC and FtsZ have so far eluded co-crystallization, probably because their complex is too unstable to be crystallized. To gain structural insight into the mechanism of action of MinC, we determined the solution structure of the N-terminal domain of *B. subtilis* MinC, and through NMR titration experiments and mutagenesis identified the binding interfaces involved in the MinC^N^-FtsZ interaction. By using our experimental results as restraints in docking, we also constructed a molecular model for the FtsZ:MinC^N^ complex and validated it by molecular dynamics. The model shows that MinC^N^ binding overlaps with the FtsZ polymerization interface on the C-terminal globular subdomain of FtsZ and, thus, provides a structural basis for MinC^N^ inhibition of FtsZ filament formation. Given that the C-terminal polymerization interface of FtsZ corresponds to the plus end of FtsZ filaments, we propose that capping is the main mechanism employed by MinC to antagonize FtsZ polymerization.

## Introduction

The Min system is a key spatial regulator of cell division in rod-shaped bacteria (Lutkenhaus, 2007; Rowlett and Margolin, 2015). It acts in concert with nucleoid occlusion proteins (SlmA in *E. coli* and Noc in *B. subtilis*) to restrict polymerization of the tubulin-homolog FtsZ to the proper midcell site, where a higher order assembly of FtsZ filaments known as the Z-ring will seed the formation of the bacterial divisome (Du and Lutkenhaus, 2019; McQuillen and Xiao, 2020). Midcell division is enforced because nucleoid occlusion proteins inhibit Z-ring formation over the nucleoids, whereas the Min system prevents division from happening at the nucleoid-free regions close to cell poles (Bramkamp and Van Baarle, 2009; Adams et al., 2014; Rowlett and Margolin, 2015). In the absence of Min, FtsZ rings will assemble at the cell poles and bacteria will divide asymmetrically producing anucleated minicells (de Boer et al., 1989; Levin et al., 1998).

In *E. coli*, the Min system consists of three proteins, MinC, MinD and MinE, which work together to generate a self-organizing oscillating protein pattern whose effect is to localize an inhibitor of Z-ring formation preferentially at the cell poles (Lutkenhaus, 2007; Rowlett and Margolin, 2015). MinC is the component of the system that interacts with FtsZ and inhibits Z ring formation and septation (Hu et al., 1999), whereas MinD and MinE are the components responsible for the oscillation (Raskin and de Boer, 1999a, 1999b). MinD is a ParA-family ATPase which oligomerizes and associates with the membrane when bound to ATP and MinE sets oscillation in motion by stimulating MinD ATP hydrolysis and membrane release (Wettmann and Kruse, 2018; Ramm et al., 2019). In addition to generating a time-averaged enrichment of MinC at the cell poles, the interaction of MinC with MinDE is important to concentrate MinC on the membrane, where Z-ring assembly takes place. Oscillating Min systems have also been described for other Gram-negative species (*Vibrio*, *Xhantomonas*) (Galli et al., 2016; Lorenzoni et al., 2017) and at least one cyanobacteria (MacCready et al., 2017). In all cases, oscillation is associated with the existence of MinE.

An alternative paradigm to the oscillating Min system is found in *B. subtilis* and related rod-shaped Gram-positives. In these bacteria, the core components MinC and MinD have been conserved and execute the same functions, but they lack MinE and MinCD instead of oscillating is stably localized to the cell poles through an indirect association with DivIVA, a widely conserved pole-marking protein of Gram-positive bacteria (Marston et al., 1998; Strahl and Hamoen, 2012; Hammond et al., 2019). MinJ is the protein that connects MinCD to DivIVA (Bramkamp et al., 2008; Patrick and Kearns, 2008), and the interplay between these four components generates a fixed gradient where MinCD concentrations are highest at the poles and decrease towards midcell, allowing FtsZ to polymerize there. Once a septation event occurs, DivIVA starts to accumulate at the future new pole by recognizing the negative curvature of the invaginating membrane and brings along MinCD, thus propagating the polar localization of these proteins. Because recruitment of MinCD to the new pole happens before cytokinesis is complete, the Z ring and MinCD proteins colocalize during the late part of the cell cycle in *B. subtilis*. Presumably, at this late stage in cytokinesis the Z ring is no longer inhibited by MinCD. Instead, evidence suggests that MinCD is necessary at this stage for the complete disassembly of Z rings upon completion of septation (Gregory et al., 2008; Van Baarle and Bramkamp, 2010; Yu et al., 2020). Z-ring disassembly has been proposed to be the main way in which MinCD prevent mini-cell formation and enforce medial division in *B. subtilis* (Gregory et al., 2008; Van Baarle and Bramkamp, 2010; Yu et al., 2020).

Despite extensive studies, the molecular mechanism used by MinC to inhibit (or disassemble) Z-rings remains incompletely understood. MinC has two clearly independent domains connected by a flexible linker (Cordell et al., 2001). The N-terminal domain (MinC^N^) of *E. coli* MinC was initially implicated as the FtsZ-inhibitory part of the protein (Hu and Lutkenhaus, 2000; Dajkovic et al., 2008a), whereas the C-terminal domain (MinC^C^) mediates MinC homodimerization and interaction with MinD (Hu and Lutkenhaus, 2000; Szeto et al., 2001). Subsequently, it was recognized that MinC^C^ is also capable of antagonizing Z-ring formation in vivo (Shiomi and Margolin, 2007; Dajkovic et al., 2008a) and genetic analysis revealed that each MinC domain binds to a different portion of FtsZ. MinC^N^ interacts with the H10 helix region at the C-terminal globular half of FtsZ, which constitutes part of FtsŹs polymerization interface, whereas MinC^C^ recognizes the conserved 15 residue patch in the unstructured tail - the “Conserved C-terminal peptide” or CCTP - of FtsZ (Shen & Lutkenhaus, 2009; Shen & Lutkenhaus, 2010). This has led to the “two-pronged model” of MinC action, in which MinC initially associates with FtsZ filaments via an interaction between MinC^C^ and the CCTP of FtsZ and this positions the MinC^N^ domain to interact with its target surface (the H10 helix) on a subunit of FtsZ in the filament (Shen and Lutkenhaus, 2009, 2010; Park et al., 2018). Because the H10 helix is part of the FtsZ polymer interface, MinC^N^ insertion between FtsZ subunits would be the ultimate cause of filament breakage. Similar, albeit less extensive, genetic data obtained with *B. subtilis* suggests that the two-pronged mechanism applies to this organism as well (de Oliveira et al., 2010; Blasios et al., 2013).

In vitro experiments also produced important insights into MinĆs mechanism. Initial experiments showed that MinC was capable of inhibiting the assembly of FtsZ polymers, but could not determine if the effect was due to filament shortening/breakage or suppression of filament bundling (Hu et al., 1999). Importantly, MinC inhibited FtsZ assembly without affecting its GTPase activity, a property that ruled out sequestration, the mechanism employed by SulA, the SOS-induced FtsZ inhibitor of gammaproteobacteria (Hu et al., 1999; Dajkovic et al., 2008b; Chen et al., 2012). Subsequent work showed that MinC (and specifically its MinC^N^ domain) can indeed shorten FtsZ filaments and established other general features of its inhibitory mechanism, such as the relatively weak affinity between FtsZ and MinC (Kd ∼ 1-10 μM), the inability to disassemble FtsZ filaments stabilized by non-hydrolyzable GTP analogs and its sensitivity to salt (Dajkovic et al., 2008a; Scheffers, 2008; Shen and Lutkenhaus, 2009, 2010; Blasios et al., 2013; Hernández-Rocamora et al., 2013; Arumugam et al., 2014; LaBreck et al., 2019). Nevertheless, some of the results obtained by the different groups are not easily reconcilable and it is still not clear how exactly MinC shortens FtsZ filaments. Examples of contradictory observations are the works from Hernández-Rocamora and collaborators (Hernández-Rocamora et al., 2013) and Arumugam and collaborators (Arumugam et al., 2014). The first described that MinC produced a homogeneous reduction of FtsZ filament size and that the protein bound preferentially to monomeric FtsZ-GDP. These observations are not compatible with a severing mechanism in which MinC would attach to filaments and break them at random positions. In contrast, Arumugam showed that MinC bound to filaments both internally and at their growing (plus) end, which corresponds to the FtsZ C-terminal polymerization face (Du et al., 2018), and caused filament shrinking by a combination of capping and increasing the off-rate of internal subunits. Whereas some of these discrepancies may be explained by the different conditions used in the experiments (Hernández-Rocamora in solution and Arumugan in a reconstituted membrane system), it is clear that more experiments are needed before we can conclude how MinC works.

The crystal structure of FtsZ in complex with other negative regulators (SulA, MciZ) has provided important structural information to corroborate in vitro experiments and establish their mechanism of Z-ring antagonism (Cordell et al., 2003; Bisson-Filho et al., 2015). However, MinC and FtsZ have so far eluded co-crystallization, probably because their complex is too unstable to be crystallized. To gain structural insight into the mechanism of MinC, we determined the solution structure of the N-terminal domain of *B. subtilis* MinC, and through NMR titration experiments and mutagenesis identified the binding interfaces involved in the MinC^N^-FtsZ interaction. By using our experimental results as restraints in docking, we also constructed a model for the MinC^N^-FtsZ complex and validated it by molecular dynamics. The model shows that MinC^N^ binding overlaps with the C-terminal FtsZ polymerization interface (the plus end) and, thus, provides a structural basis for MinC^N^ inhibition of FtsZ filament formation.

## Methods

### Expression and purification of FtsZ and MinC

DNA coding for *B. subtilis* FtsZ^1-315,A182E^ and MinC^1-102^ (hereafter referred to as MinC^N^) were subcloned into RP1B vector to generate N-terminal His-tagged proteins (plasmids and strains used in this work are listed in **Tables S1 and S2**). Site directed mutagenesis was used to create variants of MinC. For protein expression, plasmid DNAs were transformed into *E. coli* BL21(DE3) cells. Cells were grown in Luria broth in the presence of selective antibiotics at 37°C to an Abs_600nm_ of ∼ 0.8, and expression was induced by the addition of 0.5 - 1 mM isopropyl thio-β-D-galactoside. Induction proceeded overnight at 18°C prior to harvesting by centrifugation at 7,647 x *g* (15 min, 4°C). Cell pellets were stored at -80°C until purification. For NMR measurements, expression of uniformly ^2^H,^15^N or ^2^H,^15^N,^13^C-labeled FtsZ was accomplished by growing cells in D_2_O-based M9 minimal medium. ^1^H,^15^N or ^1^H,^15^N-labeled MinC was obtained by growing cells in M9 minimal medium. Both media contained 1 g/L of ^15^NH_4_Cl and /or 3 g/L of D-[^2^H,^13^C]-glucose (CIL) as the sole nitrogen and carbon sources, respectively. Multiple rounds (25, 50, 70, and 100 % of D_2_O) of adaptation were necessary for high yield expression of FtsZ.

Cells pellets were resuspended in Buffer A (50 mM Tris-HCl, pH 8.0, 500 mM KCl, 5 mM imidazole, 5 % glycerol) with 0.1 % Triton X-100, 1 mM PMSF and 100 µg/mL lysozyme for both proteins, and 20 µM GDP for FtsZ. After resuspension, cells were lysed using high pressure homogenization (Avestin C3 EmulsiFlex). The lysate was cleared by centrifugation (40,905 x *g,* 45 min, 4°C). The supernatant was filtered and loaded onto a His-Trap 5 mL column equilibrated in Buffer A. Protein was eluted in a step gradient of Buffer B (50 mM Tris-HCl, pH 8.0, 500 mM KCl, 500 mM imidazole, 5 % glycerol). Protein came out in the fraction of 30 % of Buffer B. The pooled eluted protein was incubated with tobacco etch virus (TEV) overnight at 4°C in dialysis buffer C (50 mM Tris-HCl, pH 8.0, 500 mM KCl, 5 % glycerol). The next day, a subtraction His-Trap purification was performed to remove TEV and the cleaved His tag. Final purification was achieved using SEC (Superdex 75 16/60, GE Healthcare) equilibrated in Buffer D (20 mM Hepes, pH 7.4, 500 mM KCl, 1 mM EDTA). The eluted protein was dialyzed against Analysis Buffer (20 mM Hepes, pH 7.4, 150 mM KCl, 1 mM EDTA).

### MinC^N^ NMR assignment and structure determination

^15^N-^13^C-labeled MinC^N^ was expressed from plasmid pAT6 as a 6-His-tagged protein. Cells were grown in M9 minimal medium at 37°C with 1 g/L of ^15^NH_4_Cl and 2 g/L of ^13^C glucose. The MinC^N^ protein expression was induced at OD_600_= 0.8 using 0.5 mM IPTG, after induction cells were incubated at 25°C for 8h. Following collection by centrifugation, cells were resuspended in lysis buffer (50 mM sodium phosphate buffer pH 7.4 and 50 mM KCl) and lysed by sonication. The supernatant of the cell lysate was loaded onto a HisTrap column (5 mL-GE Healthcare) and the bound protein was eluted with an imidazole gradient (0-1 M). The fractions containing MinC^N^ were collected, concentrated and cleaved with thrombin (1 U/mL) for 16h at 4°C. After cleavage, the protein was purified by size exclusion chromatography using a Sephadex 200 16/60 (GE Healthcare) column with the lysis buffer.

NMR spectra were collected from ^1^H,^15^N,^13^C-labeled MinC^N^ at 450 μM in 50 mM KCl and 50 mM Na_2_HPO_4_, pH 7.4 (10% D_2_O/90% H_2_O). All data were acquired at 293K on an Agilent 600MHz spectrometer, equipped with an inverse detection triple resonance cryoprobe, processed with NMRPipe (Delaglio et al., 1995) and visualized with NMRView (Johnson and Blevins, 1994).

To solve the solution structure of MinC^N^ the sequential backbone resonance assignments were obtained using 2D ^15^N-HSQC, 3D CBCA(CO)NH/HNCACB and 3D HNCO/HN(CA)CO experiments. Side chains for both aliphatic and/or aromatics were assigned using 3D HCCH- and CCH-TOCSY, ^13^C NOESY-HSQC and ^13^C HSQC experiments. Distance restraints for structure calculations were derived from cross-peaks in ^15^N-edited NOESY-HSQC (τm = 80 ms), ^13^C-edited aliphatic and aromatic NOESY-HSQC in H_2_O (τm = 80 ms) respectively. Peak picking was performed manually using NMRView (Johnson and Blevins, 1994).

NOE assignment and structure calculations were performed using CYANA (version 2.1) (Güntert, 2004) in a semi-automated interactive manner, using 100 starting conformers. CYANA protocol was applied to calibrate and assign NOE cross peaks. After the first rounds of automatic calculations, the NOESY spectra was analyzed again to identify additional cross peaks consistent with the structural model and to correct misidentified NOEs. Slowly exchanging amides were identified by lyophilizing the protein from water and then dissolving it in 100% D_2_O. The final 20 lowest-energy structures were refined with the CNS package. Resulting structures were analyzed using PROCHEK validation software (Laskowski et al., 1996). The final refined ensemble of 20 structures and resonance assignments for MinC^N^ domain were deposited into the Protein Data Bank (PDB ID 2M4I) and BioMagRes DB (BMRB accession number 19007), respectively.

### NMR analysis of FtsZ-MinC interaction and MinC_N_ dynamics

NMR data were collected using Bruker Avance III 800 MHz spectrometer equipped with a TCI Z-gradient cryoprobe at 298 K. NMR measurements of FtsZ or MinC were recorded using ^2^H,^15^N-labeled FtsZ or ^1^H,^15^N-labeled MinC at a final concentration of 0.1 mM in presence of ligands, in Analysis Buffer and 90 % H_2_O, 10 % D_2_O. Chemical shift perturbation (Δδ) of ^2^H,^15^N-FtsZ in the presence of unlabeled MinC or ^1^H,^15^N-MinC in presence of unlabeled FtsZ, were calculated using the following equation (Eq. 1):

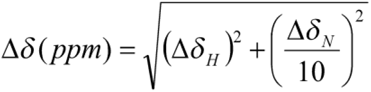

The backbone assignment of FtsZ was obtained using ^2^H,^15^N,^13^C-labeled FtsZ (0.2 mM) and through the analysis of 3D TROSY-HNCA, 3D TROSY-HN(CO)CA and 3D TROSY-HNCACB experiments. All NMR data were processed using NMRpipe (Delaglio et al., 1995) and analyzed using CcpNMR version 2.4 (Vranken et al., 2005) or CARA (http://www.nmr.ch).

MinC^N^ backbone ^15^N longitudinal (R_1_) and transverse (R_2_) relaxation rates were obtained by using inversion recovery (R_1_) and CPMG (R_2_) HSQC based Bruker pulse sequences. T_1_ and T_2_ were acquired with a recycle delay of 3 s between experiments and the following relaxation delays for T_1_: 150, 300, 450, 600, 900, 1250, 1500, 1800, 2000 and 2225 ms; and T_2_: 16.32, 32.64, 48.96, 65.28, 81.60, 97.92, 114.24, 130.56 and 163.20 ms. Systematic error in both T_1_ and T_2_ experiments were estimated from the variance of repetition experiments, which were acquired with delays of 600 ms for T_1_ and 81.60 ms for T_2_. ^1^H,^15^N-NOE (hetNOE) values were determined from a pair of interleaved spectra acquired with or without pre-saturation and a recycle delay of 5 s. Fittings of the intensity decay curves to calculate R_1_ and R_2_ relaxation rates and the hetNOE were carried out using CCPN software (Vranken et al., 2005).

### Light Scattering

For light scattering experiments, FtsZ^1-382^ and MinC^1-224^ WT and mutants were used. pET28a-FtsZ^1-382^ (pAB20) and pET24b-MinC^1-224^ (pAT30) WT and mutants were expressed in *E. coli* Bl21(DE3)-RIL (Stratagene) upon induction by 0.1 mM IPTG. FtsZ^1-382^ was purified by two-step precipitation: 30 % ammonium sulfate, followed by addition of 1 mM GTP and 20 mM Ca^2+^. MinC^1-224^ WT and mutants were purified by nickel affinity chromatography. His-MinC were eluted from the column applying an imidazole gradient (100 mM to 1 M) in HMK buffer (50 mM Hepes, pH 7.7, 5 mM Mg(CH_3_COO)_2_, 100 mM KCH_3_COO).

Light scattering by FtsZ polymers was measured using a Hitachi F-4500 fluorimeter. Excitation and emission wavelengths were set to 350 nm, with slit widths of 3.5 nm, and the photomultiplier tube at 950V with a scan rate of 60 nm/s. Protein was incubated in 150 µL of HMK buffer at 25 °C until baseline stabilization. GTP was added to 2 mM to start polymerization and the change in scattering was followed for the next 30 minutes. MinC was added to reactions either before or after the addition of GTP with similar results. Buffer effects were excluded by inclusion of MinC storage buffer to a similar volume as the maximum amount of MinC added.

### Fluorescence Microscopy

Microscopy was performed on a Nikon Eclipse Ti microscope, equipped with GFP BrightLine and mCherry BrightLineFilter Sets (Semrock), a Plan APO VC Nikon 100X objective (NA=1.4), a 25 mm SmartShutter and Andor EMCCD i-Xon camera. Exposure times varied from 0.3 to 1 s. Cells were grown to exponential phase and incubated in chambers with LB plus 1% agarose. Membrane stain FM5-95 (final concentration of 5 μg/mL; Molecular Probes) and 0.5 mM IPTG were added to the solidified LB, where indicated. All images were captured using NIS software version 3.07 (Nikon) and processed with ImageJ software (http://rsb.info.nih.gov/ij/).

### Western blotting

Cell lysates were resolved on SDS PAGE gels, transferred to PVDF membrane and probed with anti-GFP rabbit polyclonal serum (kind gift of D. Rudner) diluted to 1:3000. After washing, secondary anti-IgG conjugated with HRP (Pierce) was added at 1:10000. Immunoblots were revealed with ECL Prime Western Blotting Detection Reagent (GE Healthcare) in High Performance Chemiluminescence Film (GE Healthcare).

### Molecular docking and modeling

Protein-protein docking employed the ClusPro 2.0 web server (Kozakov et al., 2017). *B. subtilis* FtsZ and MinC^N^ structures were obtained from PDB with IDs 2VAM and 2M4I, respectively. To avoid bias in the rigid-body docking approach, we removed the flexible unstructured N and C-terminal tails of MinC^N^, (residues 1 to 10 and 94 to 105, respectively) from the structure prior to docking. Protein-protein contact information obtained from NMR and mutagenesis experiments were used to optimize docking solutions. Attraction constraints were applied to FtsZ residues 255, 256, 257, 258, 284, 285, 286, 287, 288, 289, 290 and 291 and MinC^N^ residues 12 and 15. The top 10 ranked structures provided by ClusPro were further analyzed and the solutions that caused obstruction of the FtsZ GTP binding site or the FtsZ unstructured C-terminal tail, which is not present in crystallographic structure, were discarded. Two FtsZ:MinC^N^ models that successfully met the structural criteria were further evaluated. Three other conditions were also alternatively applied in the docking approach: FtsZ:MinC^N^ docking using only MinC^N^ attraction, only FtsZ attraction or no constraints for both proteins.

In addition to an algorithm based on physical energy functions, such as ClusPro, we also modeled the FtsZ:MinC^N^ complex by Alphafold-Multimer (Evans et al., 2021). AlphaFold-Multimer (v2.1.1) was performed with a local installation from (https://github.com/deepmind/alphafold). The FtsZ and MinC FASTA sequences were used as input and obtained from PDB (PDB ID: 2VAM and 2M4I, respectively). AlphaFold was executed in multimer mode with a prokaryote flag. The maximum template data was set to 2021-01-27. The analysis and comparisons with docking models were performed using the best ranked relaxed predicted structure generated by the algorithm.

To assess which model better described the FtsZ:MinC^N^ complex interface, we compared the predicted interface with the interface identified by NMR. For this, we calculated an interface coverage as the ratio of the number of MinC^N^ or FtsZ residues from the predicted interface that presented chemical shift perturbation in NMR by the total of residues from the predicted interface, as follows (Eq. 2):

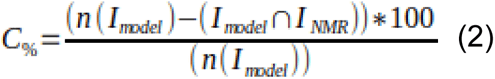

where C% is the interface coverage (0 ≥ C% ≥ 100), I_model_ is the list of FtsZ or MinC^N^ residues in the model interface, I_NMR_ is the list of residues that presented chemical shift perturbation in NMR and n(I_model_) is the length of I_model_ list.

The cut-off value used to consider that the residue had a significant chemical shift perturbation was the mean of the values of chemical shift perturbation of all analyzed residues. We also determined an interface error as the ratio of the number of MinC^N^ or FtsZ residues from the predicted interface that did not show chemical shift perturbation by the total of residues from the predicted interface, as follows (Eq. 3):

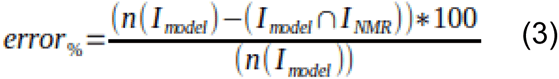

We used a distance cut-off of 5 Å, including hydrogen atoms, to define interface residues and considered only the residues that were common to the ClusPro and Alphafold-Multimer models. The best model was the one with the highest ratio between coverage and error percentage, according to equations 2 and 3.

### Molecular dynamics simulations

Molecular dynamics (MD) simulations of the selected FtsZ:MinC^N^ model were performed to evaluate the stability of the complex and the interface contacts. The simulations used the YASARA software (Krieger et al., 2002) with AMBER14 force field. Prior to simulations, we restored the MinC^N^ N- and C-terminal tails that were previously removed from the NMR structure and added ACE and NME caps to FtsZ N- and C-terminus. The FtsZ:MinC^N^ complex was solvated with water in a 15 Å cubic cell with periodic boundaries, neutralized and submitted to energy minimization. Hydrogen-containing bonds were constrained using the LINCS algorithm. Long-range electrostatic interactions were calculated using the Particle Mesh Ewald algorithm and short-range non-bonded interactions were calculated with a 8 Å cutoff. The production run was performed at 298 K and pH 7.4 with a multiple 2*2.5 fs timestep. Simulation snapshots were saved each 250 ps.

The stability of MinC^N^ in relation to FtsZ was assessed by first aligning the simulation trajectory by FtsZ and then calculating the root-mean-square deviation (RMSD) of the MinC^N^ backbone, not considering the N- and C-terminal tails, using VMD software [Humphrey et al., 1996]. The last 200 ns of the simulation, in which the system achieved RMSD convergence, was used for further analysis. Putative hydrogen bonds between FtsZ and MinC^N^ during the simulation were computed in VMD, using default criteria of donor-acceptor distance of 3 Å and angle cut-off of 20°. The trajectory was submitted to a hierarchical cluster analysis and the structure most close (minimum RMSD) to the average structure of the most populated cluster was selected as a representative model of the simulation. This structure was further refined by energy minimization.

### Data and Materials Availability

FtsZ backbone assignments have been deposited in BMRB under accession number 52358. The 3D structural models presented in this study are available in https://github.com/LBC-LNBio/MinC-FtsZ_models.

## Results

### NMR solution structure of MinC N-terminal domain (MinC^N^)

To investigate the interaction between FtsZ and MinC in atomic detail, we first solved the structure of the N-terminal domain of *B. subtilis* MinC (MinC^N^) using standard heteronuclear solution NMR techniques. We assigned 94% of MinC^N^ (96 out of 102 residues), with the missing residues being V1, K2, K15, I91, T92 and E102. An ensemble of the 20 lowest energy conformers was selected for analysis, and the structure calculation summary is given in **Table 1**. The ^1^H, ^13^C and ^15^N chemical shift assignments have been deposited in the BioMagResBank database (http://www.bmrb.wisc.edu), accession number 19007 and have been published (Castellen et al., 2015). The calculated structure was deposited in the Protein Data Bank with the accession code 2M4I. *B. subtilis* MinC^N^ shows the same overall fold as the N-terminal domain of full-length MinC from *T. maritima* (PDB 1HF2), being comprised of a four-stranded β-sheet packed against two α-helices in an approximately parallel orientation (**Figure 1**). The first eleven and the last thirteen residues are located in disordered regions, with the latter corresponding to the linker connecting MinC N-terminal and C-terminal domains (residues 1-8 and 92-102, respectively in *B. subtilis* MinC). Comparison of *B. subtilis* MinC^N^ with the crystal structures of *S. typhimurium* (PDB 3GHF) and *E. coli* MinC^N^ (PDB 4L1C) shows more significant differences, such as the presence of a third α-helix in enteric bacteria MinC^N^ and the fact that these proteins crystallized as swapped dimers (An et al., 2013). The monomeric solution structure of *B. subtilis* MinC^N^ is well supported by the protein molecular tumbling time (Τc) of 8.35 ns and by size exclusion chromatography results (not shown), and corroborates the suggestion that the swapped dimers of enteric bacteria MinC^N^ are crystallographic artifacts (Park et al., 2018).

**Figure 1:**
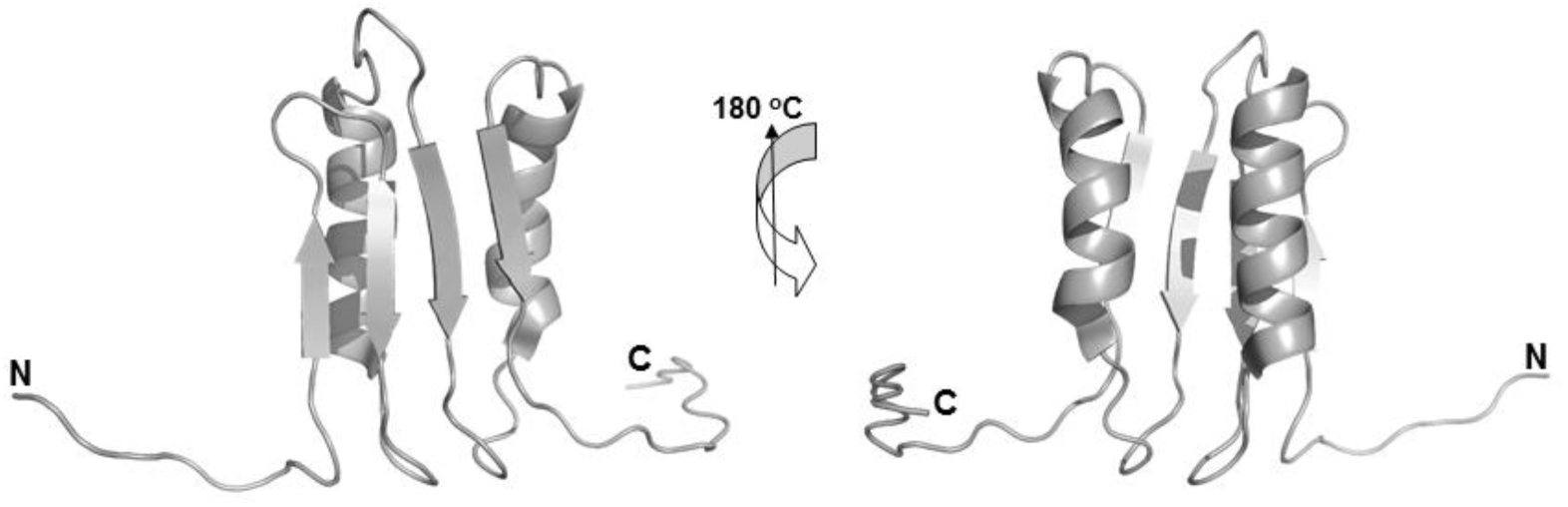
Ribbon diagram representation of the MinC^1-102^ (MinC^N^) structure from Bacillus subtilis determined by solution NMR, with the two views related by 180° rotation.

**Table 1.** NMR-based restraints used for MinC^N^ structure calculation and structure refinement statistics.

| Distance restraints |  |
| --- | --- |
| NOE | 1257 |
| Intraresidue (i,i) | 399 |
| Sequential (i, i+1) | 416 |
| Medium range (i, i+2 a i+4) | 235 |
| Long range (i, i≥5) | 207 |
| Rmsd from the Average Structure |  |
| Structured region residues, 11-90 |  |
| Backbone (structured) | 0.58±0.13 |
| Heavy atoms (structured) | 0.82±0.19 |
| Backbone (total) | 2.80±1.07 |
| Heavy atoms (total) | 3.97±1.46 |
| Structure quality |  |
| Ramachandran plot (PROCHECK) |  |
| Most favored regions (%) | 68.2 |
| Additional allowed regions (%) | 31.3 |
| Generously allowed regions (%) | 0.5 |
| Disallowed regions (%) | 0.0 |
| CYANA |  |
| Target function (Å) | 0.24±0.08 |
| Distance violation >0,20 | 0 |
| Dihedral angles >5Å | 0 |

To further verify our structure, we performed auto-correlated fast timescale ^15^N relaxation experiments and observed that MinC^N^ exhibits highly uniform ^15^N dynamics throughout its backbone with average R_1_ of 1.03 ± 0.01 s^-1^, R_2_ of 14.98 ± 0.7 s^-1^ and hetNOE of 0.7 ± 0.01 (**Supplemental Figure 1**). Increased dynamics are consistent with the structure, being restricted to the expected flexible regions, such as loops and in the disordered N- and C-termini. Interestingly, MinC^N^ also showed regions - the end of helices H1 and H2 and the loop H2-S4 - whose measured dynamics indicates conformational exchange. Thus, MinC^N^ may show alternative conformations important for its interactions and function.

### NMR mapping of the MinC^N^ - FtsZ interaction

To investigate the binding site for FtsZ in MinC^N^, we carried out titration experiments using as the ligand *B. subtilis* FtsZ lacking its unstructured C-terminal tail and with a mutation to keep the protein monomeric at the high concentrations necessary for NMR (A182E, equivalent to A181E of *E. coli*) (Li et al., 2013). We recorded 2D [^1^H,^15^N] HSQC NMR spectra of MinC^N^ with increasing concentrations of FtsZ^1-315,A182E^ (1:0.12, 1:0.25, 1:0.5 molar ratio). Many MinC^N^ residues broadened beyond detectability upon FtsZ addition and a few residues showed chemical shift perturbations compared to the MinC^N^ apo spectrum (**Figure 2A,C**). This indicates binding in the intermediate exchange regime, which is in line with the modest affinity between MinC and FtsZ (∼1-10 μM, depending on the pH and ionic strength) (Hernández-Rocamora et al., 2013; Park et al., 2018). Mapping of the missing resonances and CSPs onto the MinC^N^ structure showed that they were found predominantly on the beta-sheet and on the upper (C-terminal) part of helices α_1_ and α_2_ and the loops connecting these elements, suggesting these regions of the protein either make direct contact or are altered upon interaction with FtsZ (**Figure 2B**). Interestingly, several of the residues perturbed in our titration experiments correspond to mutations identified by the Lutkenhaus and Camberg labs which disrupt MinC ability to antagonize FtsZ (Gly13, Thr19, His21 and Thr45, respectively Gly10, Ser16, Val18, and Phe42 in *E. coli*) (Park et al., 2018; LaBreck et al., 2019). Lys12, a highly conserved lysine next to Gly13 and which is also important for MinC function in *E. coli* (Lys9) could not be interrogated because it was not assigned in our spectrum.

**Figure 2:**
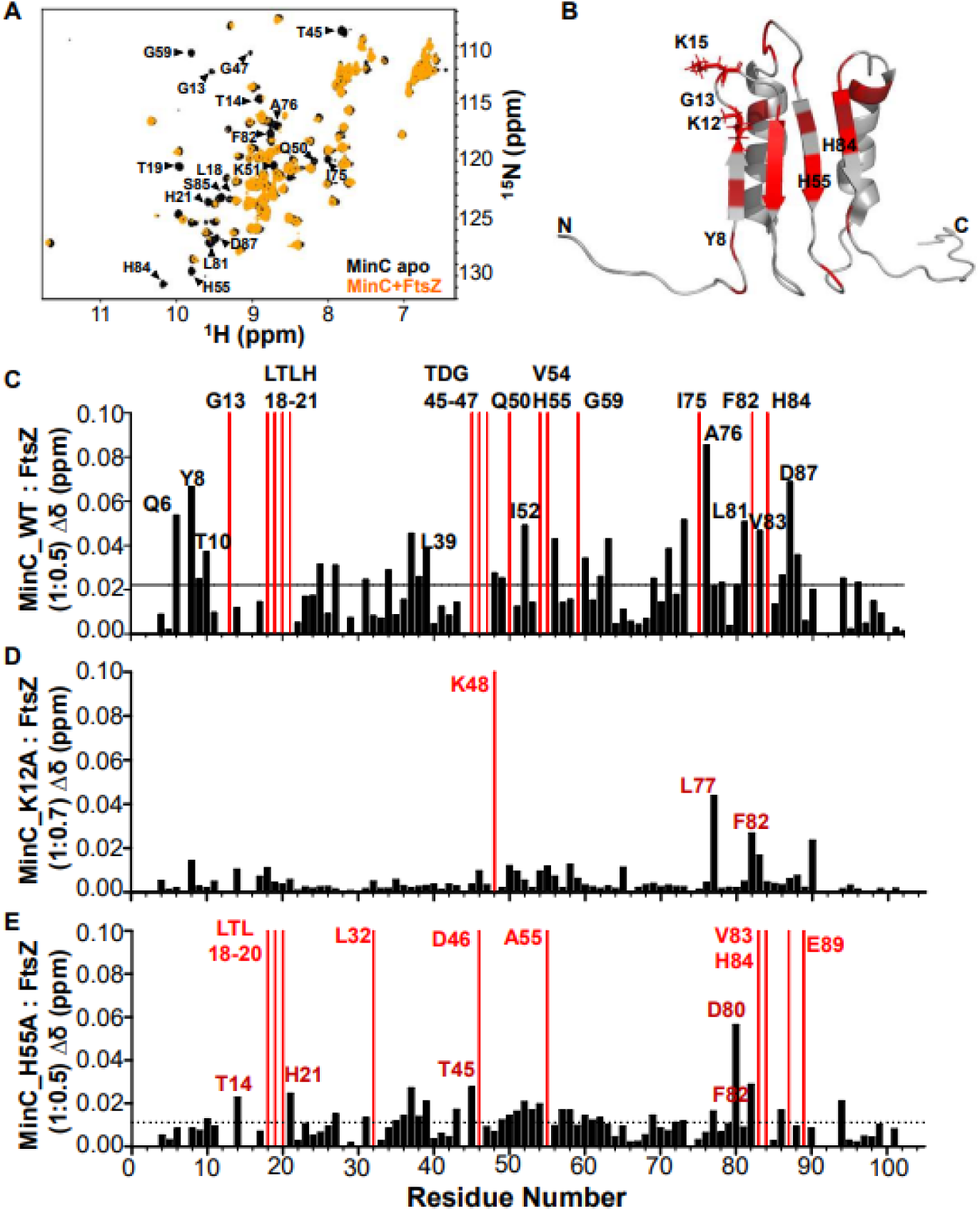
Titration of MinC^N^ with FtsZ^1-315,A182E^. (A) [^1^H,^15^N]-HSQC of MinC^N^ (100 μM) in black overlapped with MinC^N^ (100 μM) in the presence of FtsZ^1-1315,A182E^ (50 μM) in magenta. (B) Ribbon representation of MinC^N^ with CSPs highlighted in dark red, and residues broadened beyond detectability upon FtsZ binding highlighted in red. (C) CSP plot of MinC^N^ versus MinC^N^ in the presence of FtsZ (1:0.5 ratio). Residues with CSPs upon FtsZ binding (σ = 0.021 ppm) are shown as black bars and residues that broadened beyond detectability are shown as red bars. (D) CSP plot of MinC^N^ mutant K12A in response to FtsZ. (E) CSP plot of MinC^N^ mutant H55A in response to FtsZ.

Next, to determine the binding site for MinC^N^ in FtsZ, we carried out the reciprocal titration experiment by collecting 2D [^1^H,^15^N] TROSY NMR spectra of ^2^H,^15^N-FtsZ^1-315,A182E^ with increasing concentrations of MinC^N^ (1:0.25, 1:0.5, 1:1, 1:2, 1:4 molar ratio). This identified CSPs and a few residues broadened beyond detectability predominantly in the C-terminal globular subdomain of FtsZ. Within this region, the majority of the altered resonances mapped to two clusters of residues between positions 250 and 300, which correspond to helix H10 and neighboring beta strands (**Figure 3A-C**). Previous genetic and biochemical experiments from the Lutkenhaus lab and our own group had already suggested that the binding site for MinC^N^ was located in the C-terminal globular half of FtsZ (Shen and Lutkenhaus, 2009, 2010; Blasios et al., 2013). In *E. coli*, mutations conferring resistance to MinC were mostly on helix H10 (L270V, R271G, E276D, N280D, I294T), whereas in *B. subtilis* mutations mapped to a similar but broader area, encompassing loops H9-β8 and H10-β9 in addition to helix H10 (K243, I245, D255, V260, A285, D287 and V310). Comparison of NMR and mutagenesis showed a remarkable correspondence in the affected FtsZ regions, corroborating the previous prediction that the MinC binding site in FtsZ is located in the vicinity of helix H10.

**Figure 3:**
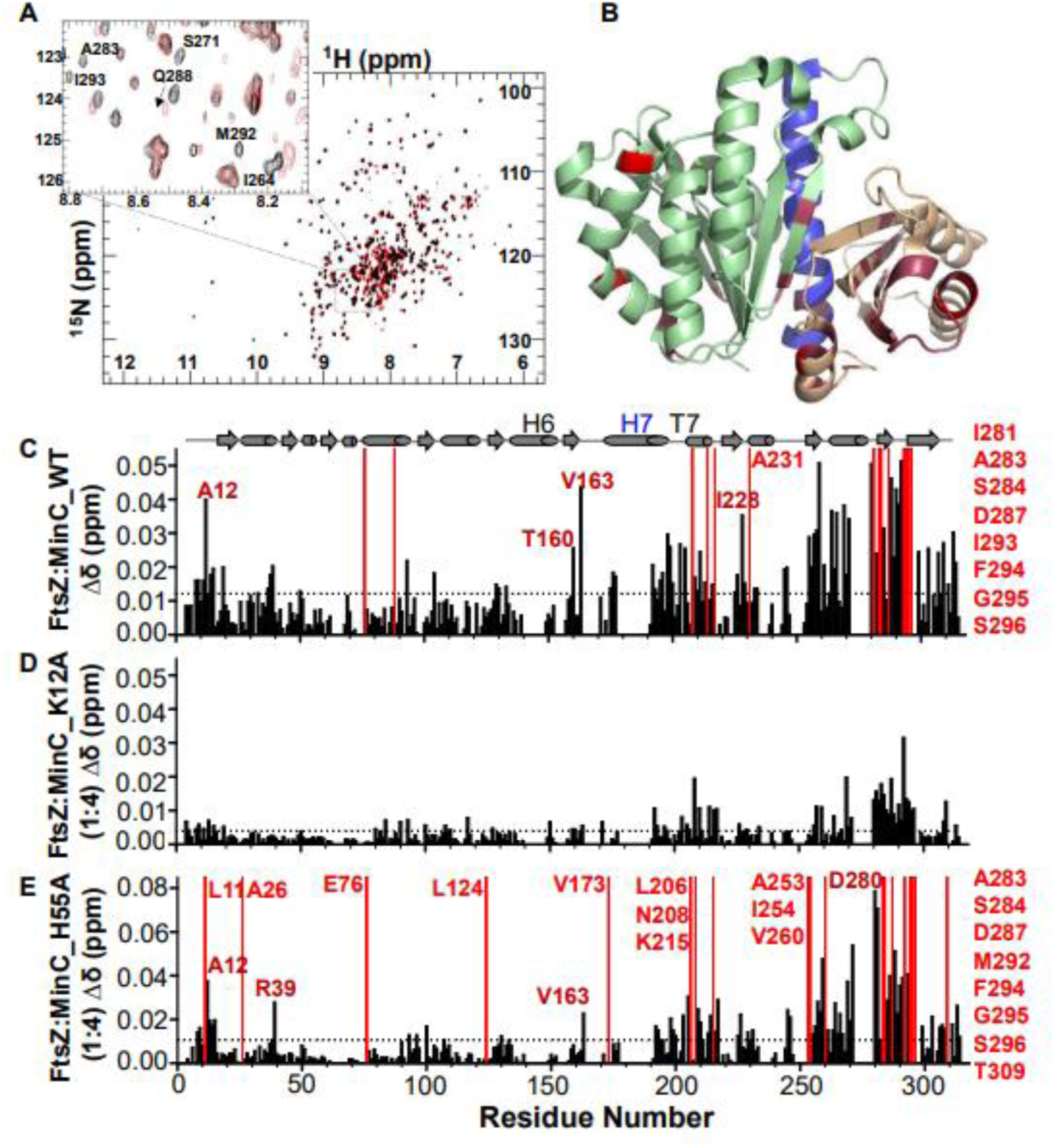
Titration of FtsZ^1-315,A182E^ with MinC^N^. (A) [^1^H,^15^N]-TROSY of FtsZ^1-315,A182E^-GDP (90 μM) in black overlapped with FtsZ^1-315,A182E^-GDP (90 μM) in the presence of MinC^N^ (360 μM) in red. (B) Ribbon representation of FtsZ [PDB:2VAM] highlighting CSPs in maroon, and residues broadened beyond detectability in red. (C) CSP plot of FtsZ^1-315,A182E^-GDP versus FtsZ^1-315,A182E^-GDP in the presence of MinC^N^ (1:4 ratio). Residues with CSPs upon FtsZ binding (σ = 0.012 ppm) are shown as black bars and residues that broadened beyond detectability are shown as red bars. (D) CSP plot of FtsZ^1-315,A182E^-GDP in response to MinC^N^ mutant K12A. (E) CSP plot of FtsZ^1-315,A182E^-GDP in response to MinC^N^ mutant H55A.

### Genetic testing of the predicted MinC^N^ interface

We tested the NMR prediction of FtsŹs binding surface on MinC^N^ by making mutants and assaying their function in vivo and in vitro. Because the N-terminal domain of MinC is not highly conserved between Gram+ and Gram- bacteria, instead of substituting the same residues that were mutated in *E. coli* MinC, we decided to choose residues to mutate based on three criteria: i) be highly conserved in Gram+ bacteria; ii) be positively charged, given that a positive charge in MinC^N^ seems crucial for FtsZ binding (Park et al., 2018; LaBreck et al., 2019), and iii) be solvent exposed and distributed over most of the interface of MinC^N^ defined by NMR. Based on these criteria we created five substitutions (Y8A, K12A, K15A, H55A and H84A) and tested their ability to complement a *minC* null mutant. Mutations were constructed in the context of a GFP-MinC fusion, to allow monitoring of mutant protein stability and localization, in addition to function.

A *minC* null mutant exhibited frequent asymmetric division and minicell formation (16%), with consequent broadening of the cell length distribution, detectable as an increase in average cell length (from around 4.5 to almost 7 μm). Expression of the wild-type GFP-MinC fusion complemented this phenotype, as seen by the restoration of cell length and minicell formation to wild-type levels. In contrast, none of the mutant GFP-MinC fully restored proper cell division, suggesting that all the mutations perturb MinC function to some extent (**Table 2**). Using minicell frequency as a measure of function, the strongest mutation was K15A (7.8%), followed by Y8A and K12A (5.1 and 4.6%), and H55A and H84A (3.7 and 2.9%) being the mildest ones. Focusing on cell length, all mutations seemed similarly defective, with the exception of Y8A, whose length was closer to the wild type (**Supplemental Figure 2A**). All mutant proteins were expressed and accumulated to similar levels (**Supplemental Figure 2C**), ruling out protein instability as the explanation for their reduced function.

**Table 2.** Cell division phenotype of MinC WT and mutants.

|  | *minC*<sup>-</sup> | WT | Y8A | K12A | K15A | H55A | H84A |
| --- | --- | --- | --- | --- | --- | --- | --- |
| Average cell size (μm) | 6.88 | 4.75 | 4.81 | 6.02 | 5.49 | 5.88 | 5.60 |
| Minicell frequency (%) | 16.1 | 1.0 | 5.1 | 4.6 | 7.8 | 3.7 | 2.9 |

In addition to binding to FtsZ, MinC needs to interact with MinD to localize and function properly. We analyzed the subcellular localization of the mutant GFP-MinC and observed that they accumulate on cell poles and nascent septa, just like the wild-type (**Supplemental Figure 2C**). Given that the interaction with MinD involves MinC^C^ and the mutations being assayed are in MinC^N^, this result is expected. Nevertheless, it indicates that the mutations do not destabilize the proteins or have pleiotropic effects, and is consistent with our hypothesis that the altered residues are specifically involved in the interaction between MinC and FtsZ.

We next analyzed the ability of MinC^N^ mutants to inhibit FtsZ polymerization in vitro, using light scattering. This time we expressed and purified wild-type and mutant MinC without a GFP fusion. In our assay condition (pH 7.7 and 100 mM potassium), wild-type MinC at a 3:1 ratio inhibited FtsZ polymerization by approximately 50%. Similar inhibition was achieved by mutants H55A and H84A, indicating that these substitutions do not significantly affect the FtsZ-MinC interaction. In contrast, MinC proteins bearing the Y8A, K12A and K15A mutations were significantly less able to inhibit FtsZ polymerization (**Figure 4**). The K12A and K15A mutants were particularly impaired, inhibiting FtsZ polymerization by just 20-25%, thus being half as active as the wild-type protein. Interestingly, the behavior of the mutants in the in vitro assay closely paralleled their in vivo minicelling phenotype: in both assays, the K12A and K15A mutations had the strongest effect, whereas H55A and H84A had the mildest effects. Together, these data suggest that not every portion of MinC^N^ perturbed in the titration experiments is equally important for binding to FtsZ. Given that CSPs do not necessarily indicate a direct interaction of the residue perturbed, it is possible that the binding interface for FtsZ may not encompass the whole beta-sheet of MinC^N^, being restricted to beta strands 1 and 2 and the connecting loop, the regions where residues K12 and K15 are located. In this case, the CSPs detected for H55 and H84 could reflect a general rearrangement of MinC^N^’s beta-sheet upon FtsZ interaction.

**Figure 4:**
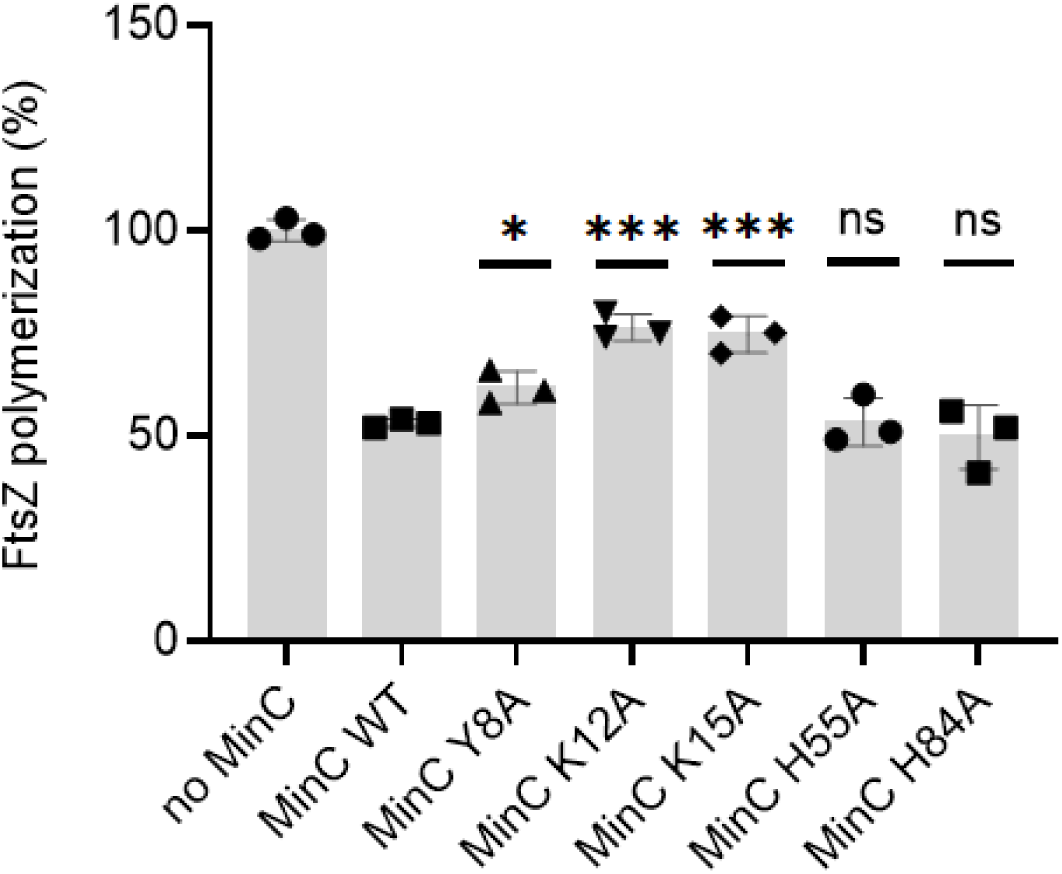
FtsZ polymerization in the presence of MinC WT and mutants measured by light scattering. Reactions contained 5 μM FtsZ in the absence (first bar) or presence of 15 μM MinC WT or mutants in HMK buffer (50 mM Hepes, pH 7.7, 5 mM magnesium acetate, 100 mM potassium acetate). MinC mutants were compared with wild-type MinC by unpaired t-tests: ns, P > 0.05; * P ≤ 0.05;*** P ≤ 0.001.

To further characterize the effects of mutations K12A and H55A, we employed NMR to directly measure binding of these mutants to FtsZ. FtsZ titration experiments showed an almost complete loss of CSPs for MinC^N^ K12A, even at a high MinC^N^-FtsZ ratio (**Figure 3D**), indicating that this substitution severely impairs the MinC^N^-FtsZ interaction. In contrast, and in line with its mild effect in vivo and in vitro, the pattern of CSPs produced by FtsZ in MinC^N^ H55A was similar to that observed in wild-type MinC_N_ (compare **Figure 3C and E**), indicating that the MinC^N^-FtsZ interaction was generally preserved in this mutant. Reciprocal experiments titrating labeled FtsZ with unlabeled MinC^N^ K12A and H55A produced similar conclusions: whereas K12A hardly perturbed the FtsZ spectrum, H55A induced similar CSPs as wild-type MinC^N^ (**Figure 4 D and E**). These results are consistent with the hypothesis that the binding site for FtsZ in MinC^N^ is centered around the loop connecting strands S1 and S2, but does not seem to include contacts with the complete beta-sheet of MinC^N^. This conclusion is also in agreement with the mutations identified in *E. coli* MinC^N^ that abolish FtsZ antagonism and which are either on the hairpin connecting strands S1 and S2 and on helix α1, but not beyond these elements (Park et al., 2018; LaBreck et al., 2019).

### Computational modeling of FtsZ:MinC^N^ complex

Our NMR analysis of the MinC-FtsZ interaction, together with extensive mutagenesis work by us and others, provided an excellent set of constraints to be applied in the computational modeling of the FtsZ:MinC^N^ complex. Protein-protein docking of FtsZ monomer and MinC^N^ was performed with the ClusPro 2.0 web server (Kozakov et al., 2017) and informed by NMR and mutagenesis experiments, using attraction and repulsion constraints. This generated two possible binding models of MinC^N^ to FtsZ (**Supplementary Figure 3A**). The docking model A was better ranked than model B, using cluster population criteria of ClusPro. Despite the different MinC^N^ orientation in relation to FtsZ in these models, almost in a mirrored configuration, both shared a similar interface involving the MinC^N^ S1 and S1-S2 turn (where K12 and K15 are located) and the FtsZ final portion of H10 and S9-H10 loop (where are located residues that showed significant CSPs in NMR, as well as that conferred resistance to MinC when mutated). To investigate whether this interface was not biased by the experimental constraints applied, we also performed the same docking protocol in three other conditions: using only MinC constraints, using only FtsZ constraints and without constraints. Importantly, all solutions produced by these alternative conditions provided an interface similar to model A (**Supplementary Figure 3B**).

We also modeled the FtsZ:MinC^N^ interface using AlphaFold-Multimer (Evans et al., 2021). Differently from the docking protocols described above that were guided by experimental data, this approach was not informed by any kind of experimental data produced in our study. The AlphaFold model shared a similar interface with docking model A, but with MinC^N^ oriented perpendicularly in comparison to model A (**Supplementary Figure 3C**). Differently from the rigid-body docking, the AlphaFold modeling allowed the detachment of MinC^N^ S1 from H1. This allowed MinC^N^ S1 and S1-S2 turn in the AlphaFold model to assume a similar orientation as observed in model A, despite their orthogonal displacement (**Supplementary Figure 3C, inset**). In both models MinC^N^ K12 and K15 point towards negatively charged residues in FtsZ H10-S9 loop and H8 (**Supplementary Figure 3C, inset**). Thus, model A and the AlphaFold model, which were obtained from different and independent computational approaches, converged to an interface region mainly involving FtsZ H10, S9, H10-S9 loop and H8.

To gain further insights on the FtsZ:MinC^N^ interface, we selected the model that best explained the CSPs and submitted this model to MD simulations without constraints. The criteria used to select the best model was the balance between the percentage of FtsZ or MinC residues that are in the model interface and are explained by NMR (“interface coverage”) and the percentage of those residues that are in the model interface but are not explained by NMR (“interface error”, see methods for details). By these criteria, the docking model A (**Figure 5A**) was the model that provided the better balance of the ratio between coverage and error for MinC and FtsZ. In addition, part of the FtsZ:MinC^N^ interface indicated by model A was also corroborated by the AlphaFold model.

**Figure 5.**
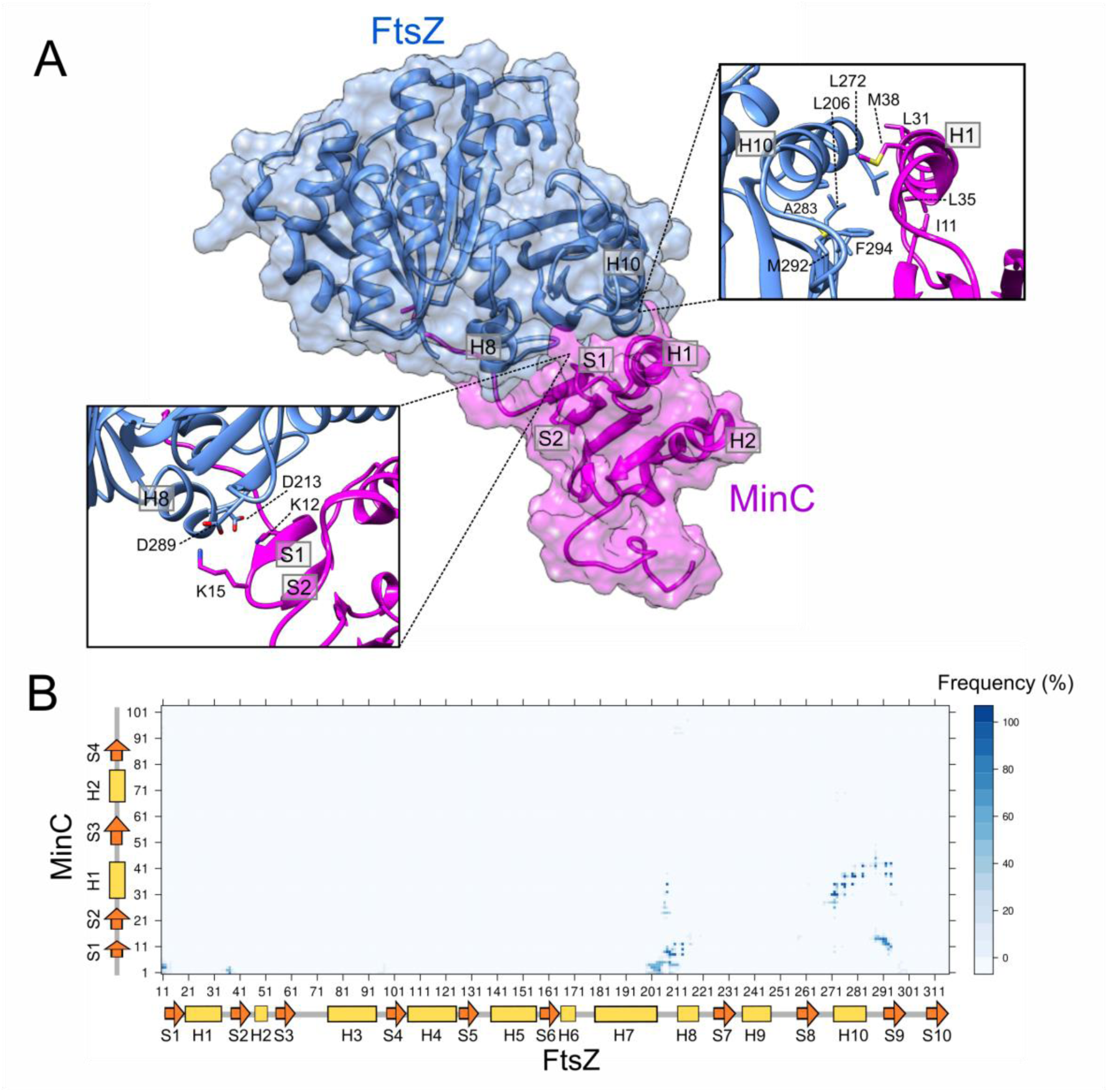
Structural model of MinC interaction with FtsZ. (A) Refined 3D structural model of the interaction between FtsZ (blue) and MinC (magenta) originated from the MD simulations of the model A. Both proteins are represented as cartoon and transparent molecular surface. Left inset shows the interaction between MinC positively charged residues K12 and K15 from S1 with FtsZ negatively charged residues of H7-H8 loop and H8. Right inset shows a helix-helix interaction between FtsZ H10 and MinC H1. Side-chains of hydrophobic residues involved in this interaction are represented as sticks. (B) Level plot of the contacts between MinC and FtsZ residues during the last 200 ns of the MD simulation of the model A. In the color scheme, dark blue (i.e. frequency of 100 %) means that the contact between the residues was observed in all simulation frames. Secondary structure of each protein is indicated in the plot to facilitate residues identification. The contacts were determined using Bio3D R package (Grant et al., 2006) with a distance cut-off of 5 Å between any atoms, considering hydrogens.

In MD simulations, the stability of the proposed FtsZ:MinC^N^ interface was evaluated by the MinC^N^ RMSD in relation to FtsZ. MinC^N^ remained stable during 400 ns of simulation with an average deviation of 5.7 Å from the initial configuration (**Supplementary Figure 4**). The contact map during the last 200 ns revealed persistent contacts of the MinC^N^ N-terminus and S1 with the FtsZ H8 and H7-H8 loop; MinC^N^ S1 with FtsZ S9; and MinC^N^ H1 with FtsZ H10 and S9 (**Figure 6B**). Putative hydrogen bonds (HB) were observed between side-chains of MinC^N^ K12 or Y8 and FtsZ D213, between side-chains of MinC^N^ K15 or K2 and FtsZ D289, and also between side-chains of MinC^N^ E42 and FtsZ Q288 (**Table S3**). The refined structural model of the FtsZ:MinC^N^ complex indicated that MinC anchors to FtsZ through two sets of contacts: electrostatic interactions between MinC^N^ K12 and K15 with FtsZ D213 and D289, respectively (**Figure 6A, left inset**), and hydrophobic helix-helix interactions between MinC^N^ H1 and FtsZ H10 (**Figure 6A, right inset**).

## Discussion

We applied NMR to study the interaction between the N-terminal domain of MinC and FtsZ and used this information together with genetic data to generate a model for the MinC^N^-FtsZ complex. We focused on the N-terminal domain of MinC because this is the part of the protein that affects FtsZ filament assembly and length (Hu and Lutkenhaus, 2000; Dajkovic et al., 2008a). Despite extensive work by several laboratories over many years, the mechanism of MinC^N^ inhibition of FtsZ polymerization is still incompletely understood and controversial (Dajkovic et al., 2008a; Scheffers, 2008; Shen and Lutkenhaus, 2010; Blasios et al., 2013; Hernández-Rocamora et al., 2013; Arumugam et al., 2014; Park et al., 2018; LaBreck et al., 2019). One important missing link is structural information showing how MinC^N^ and FtsZ interact, something that has eluded attack by crystallography or cryoEM because the proteins do not form a stable complex. However, NMR can still provide valuable structural insight about transient protein-protein interactions that cannot be captured by other methods (Clore and Venditti, 2013). Our results indicate that MinC^N^ binds to FtsZ at the bottom of its C-terminal globular domain, which corresponds to the plus end of the protein, by contacting predominantly helix H10, and strands S8 and S9 and covering a large part of the surface that FtsZ uses to interact with itself. Such binding site is in principle compatible with MinC^N^ acting by capping or severing of FtsZ filaments, or even by sequestration of FtsZ monomers, but rules out other models in which MinC^N^ effect would occur indirectly via inhibition of filament bundling, as previously proposed by ourselves (Blasios et al., 2013). Furthermore, it provides a structural basis to inform further experimentation to fully elucidate the mechanism of this prominent yet still mysterious regulator of bacterial cell division.

The experiments and model described here employed *B. subtilis* proteins, but we expect that their conclusions should be generally applicable, including to *E. coli* where most work with MinC has been carried out. Despite the low conservation of the amino acid sequence, the structure of *B. subtilis* MinC^N^ has the same topology and is quite similar to MinC^N^ of other bacteria whose experimental structures are available (*T. maritima* - 1HF2; *E. coli* - 4L1C; *S. typhymurium* - 3GHF). Moreover, mutations that perturb the MinC-FtsZ interaction identified in *E. coli* and *B. subtilis* map to similar regions and, in some cases, identical or equivalent residues in both species. Examples are the mutations in the H10 helix of FtsZ that make it resistant to MinC (de Oliveira et al., 2010; Shen and Lutkenhaus, 2010; Blasios et al., 2013), and the conserved lysine in MinĆs first beta-strand/loop (K9 in *E. coli* and K12 in *B. subtilis*), whose substitution strongly diminishes MinC^N^ inhibitory activity (Park et al., 2018; LaBreck et al., 2019) and this work). Lastly, the biochemical properties of *B subtilis* MinC, despite less thoroughly explored, are in general agreement with *E. coli*’s, including the shortening of filaments without inhibition of FtsZ GTPase, the salt sensitivity of the interaction with FtsZ and the rescuing effect of ZapA (Scheffers, 2008; Blasios et al., 2013).

Our model of the FtsZ:MinC^N^ complex is very similar to the one predicted by AlphaFold Multimer, with the same surfaces of FtsZ being used in both cases, strengthening the conclusion that the S8, S9 and H10 elements in the plus end of FtsZ are the binding site for MinC^N^. Interestingly, however, there was a difference in the orientation of MinC^N^ in the two models, with MinC^N^ in Alpha Fold’s rotated around 90 degrees compared to ours. The altered orientation also translated into different contacts made by MinC^N^, which included helix H1 in our model but not in AlphaFold’s. The reason for this disagreement between the models seems to be a difference in the structure of MinC^N^ itself, which have alternative orientations of the helices H1 and H2 relative to the beta sheet formed by S1-4, when we compare our experimental structure with AlphaFold’s. A recent survey of the accuracy of AlphaFold models showed that even very high-confidence predictions can differ from experimental maps on a global scale through distortion and domain orientation, as observed here (Terwilliger et al., 2024). Moreover, the discrepant MinC^N^ regions are predicted to be of high to low- confidence in the Alpha Fold model of MinC (https://alphafold.ebi.ac.uk/entry/Q01463). To seek further evidence of which model better reflects reality, we have also evaluated both complexes by comparing their general interface features (buried area, polar vs nonpolar contacts, etc) and how the predicted interfaces match with mutations shown to disrupt the MinC^N^-FtsZ interaction in *B. subtilis* and *E. coli*. The interfaces in both complexes have similar overall features, with a slightly larger buried area and frequency of nonpolar contacts in the case of our complex (**Table S4**). Our model was also slightly better in how well the predicted interfaces matched the MinC^N^ mutant data (**Table S5**). Thus, although our analysis suggests that both complexes are generally equivalent, we would like to argue that the orientation of MinC^N^ in our complex is more likely to be correct, as it is based on an experimental structure. Irrespective of which structure is correct, they are equivalent in the way FtsZ is affected and hence the mechanistic implications will be the same, as described below.

How does the proposed structure of the FtsZ:MinC^N^ complex advances our understanding of MinĆs mechanism? A widely accepted view of how MinC shortens FtsZ filaments is the “two-pronged model”, put forth by Lutkenhaus and collaborators (Shen and Lutkenhaus, 2009, 2010). This model proposes that MinC gets recruited to FtsZ filaments via an interaction between FtsŹs CTP and MinCC and this, in turn, positions MinC^N^ so that it can break the nearest Z-Z bond. Such breaking of filaments at internal bonds is known as severing and has been shown to occur for actin and microtubules and require some form of active splitting of subunits, often by imposing mechanical stress on the filament (Ono, 2007; McNally and Roll-Mecak, 2018). Because MinC only affects FtsZ filaments that hydrolyse GTP, one assumption is that MinC^N^ can only break bonds that are occupied by GDP and which are flexible enough to allow MinC^N^ to wedge itself in between FtsZ subunits to find its binding surface. Even though this model is consistent with genetic and some biochemical data, our prediction that MinC^N^ and FtsZ share a large interaction surface (2353 Å^2^ buried area) and its location far inside the FtsZ polymerization interface suggests that it is unlikely that MinC^N^ would be able to reach its binding site by “sneaking in” between FtsZ subunits of a filament, even if these are connected via a GDP bond. Both molecular dynamics as well as structures of filaments in GDP-bound state have shown that the elements recognized by MinC^N^ in FtsZ (H10, S9, S8) remain in intimate contact with the neighboring FtsZ subunit when GDP is at the interface (Ramírez-Aportela et al., 2014; Fujita et al., 2017; Schumacher et al., 2020; Yoshizawa et al., 2020; Ruiz et al., 2022). In fact, opening of the interface upon GTP hydrolysis is at the opposite side recognized by MinC^N^ (see Ruiz et al., 2022 and MovieS2 in Ramírez-Aportela et al., 2014). Thus, given that MinC^N^ can only access its binding surface on FtsZ after the filament is broken, we conclude that for severing to occur another part of MinC (MinC^C^?) must be in charge of breaking the filament to expose MinC^N^’s binding site. However, data from several laboratories showed that MinC^C^ or full length MinC with mutations in MinC^N^ are not capable of shortening or breaking FtsZ filaments (Dajkovic et al., 2008a; Park et al., 2018; LaBreck et al., 2019). Another argument against severing is the observation that MinC-treated FtsZ filaments maintain a narrow size distribution, instead of the broader one expected if filament breakage occurred randomly along their lengths (Hernández-Rocamora et al., 2013). Thus, our structural predictions, together with previously published biochemical data, are inconsistent with MinC acting as a severing protein. Instead, the binding site for MinC^N^ seems more consistent with MinC acting by capping or sequestration. Capping the plus end of FtsZ filaments is a highly effective way to promote filament shortening by blocking addition of subunits to its growing end, as shown for MciZ, and more recently for MipZ (Bisson-Filho et al., 2015; Corbin and Erickson, 2020; Corrales-Guerrero et al., 2022). Sequestration is also an effective way to shorten filaments and the mechanism employed by SulA (Dajkovic et al., 2008b; Chen et al., 2012; Corbin and Erickson, 2020), and probably also by the less well characterized modulators OpgH (Hill et al., 2013) and Kil (Hernández-Rocamora et al., 2015), but usually requires high affinity for FtsZ, much higher than the affinity displayed by MinC. Capping and sequestration can have similar effects on filament length distribution and it is not uncommon that cappers will also display sequestration activity depending on the experimental regimen (Bisson-Filho et al., 2015). However, cappers and sequesterers differ in one critical way: whereas cappers are capable of increasing the GTPase activity of FtsZ (via increased filament ends/turnover), sequesterers always reduce FtsŹs GTPase (Corbin and Erickson, 2020). Although no systematic studies have been carried out to investigate MinC effect on FtsZ GTPase under different stoichiometries (something necessary to reveal increases in GTPase activity), the fact that MinC shortens filaments without reducing FtsŹs GTPase strongly suggests that this negative modulator functions primarily as a capper.

Another noteworthy conclusion from this work is that the binding site recognized by MinC^N^ in FtsZ overlaps substantially with the ones employed by SulA and MciZ, other negative modulators that inhibit filament formation by sequestration and capping respectively, and the only ones with a crystal structure in complex with FtsZ (Cordell et al., 2003; Bisson-Filho et al., 2015) (**Supplemental Figure 5**). In addition, recent data indicates that MipZ, a *C. crescentus* negative modulator responsible for Z ring positioning also interacts with the H10 region of FtsZ (Corrales-Guerrero et al., 2022). Thus, the polymerization surface on the C-terminal globular domain of FtsZ - the plus end - seems to be a hotspot for inhibitors of filament formation and we can speculate why this is so. One possibility is that the plus end surface of FtsZ has structural features that make it easier to accommodate new protein-protein interactions. More likely, however, inhibitors evolved to hit FtsZ on the plus end, as this is the most effective way of promoting the disassembly of filaments that have kinetic polarity and undergo treadmilling like actin and FtsZ (Bisson-Filho et al., 2015; Wagstaff et al., 2017; Corbin and Erickson, 2020). Binding to the N-terminal polymerization face - the minus end - would instead promote filament nucleation and stabilization (Mullins et al., 1998). Targeting the N-terminal face would only inhibit filament formation if that caused a concurrent conformational change that prevented that subunit from associating with a filament, i. e., a sequestration mechanism.

In conclusion, we have applied a combination of NMR and modeling to produce the first atomic description of the FtsZ-MinC interaction and the information obtained strongly suggests that MinC inhibits FtsZ polymerization by a capping mechanism. A similar strategy could be employed to advance ournowledge of other FtsZ negative regulators which are unsuitable for crystallography and whose mechanisms remain poorly understood.

---

**Supplemental Figure 1:**
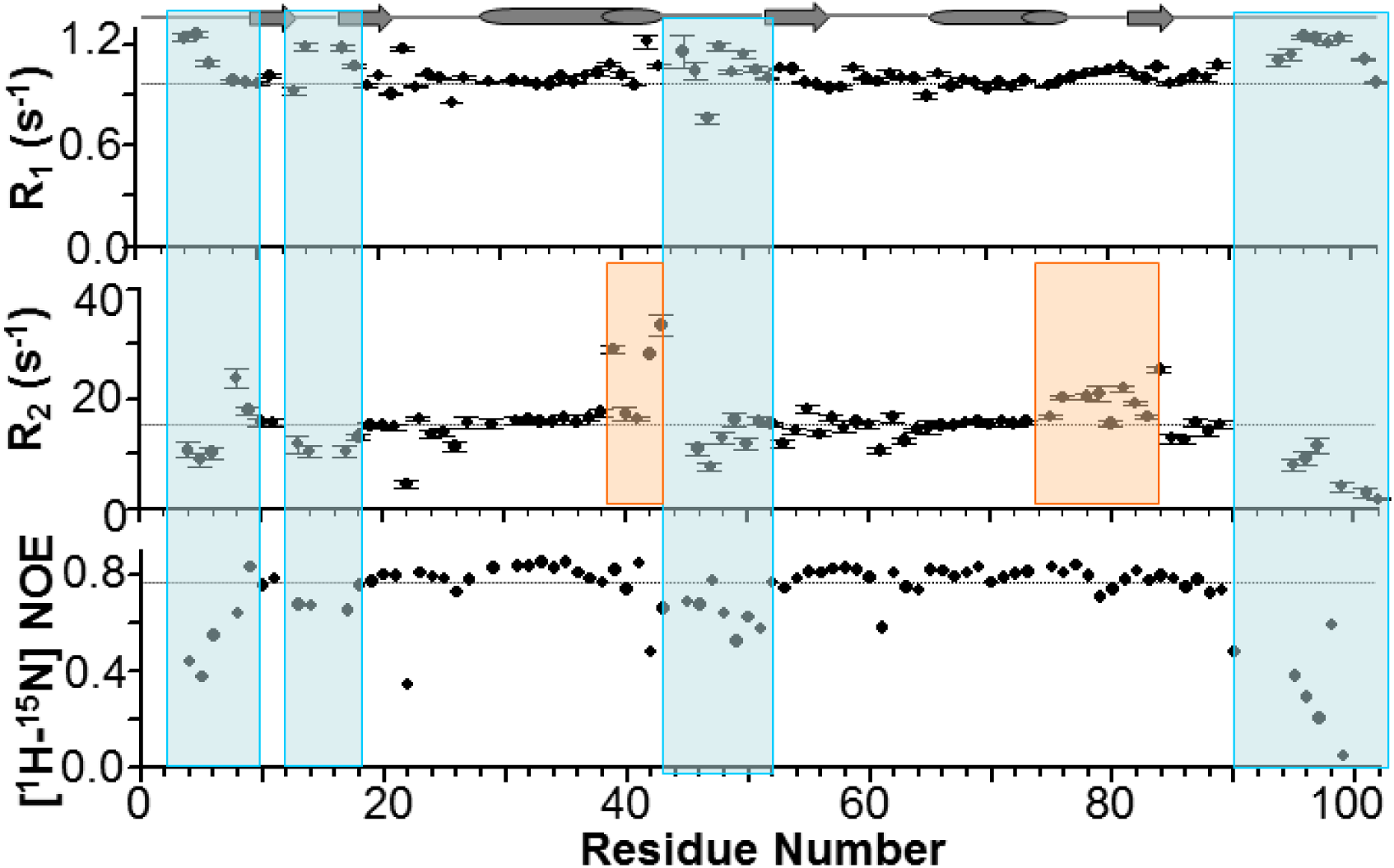
^15^N dynamics of MinC^N^. ^15^N R_1_ and R_2_ relaxation rate and hetNOE (^15^N 80 MHz Larmor frequency). Annotated secondary structure is shown. The dynamic loops are highlighted in blue. Regions of potential conformational exchange are highlighted in orange.

**Supplemental Figure 2:**
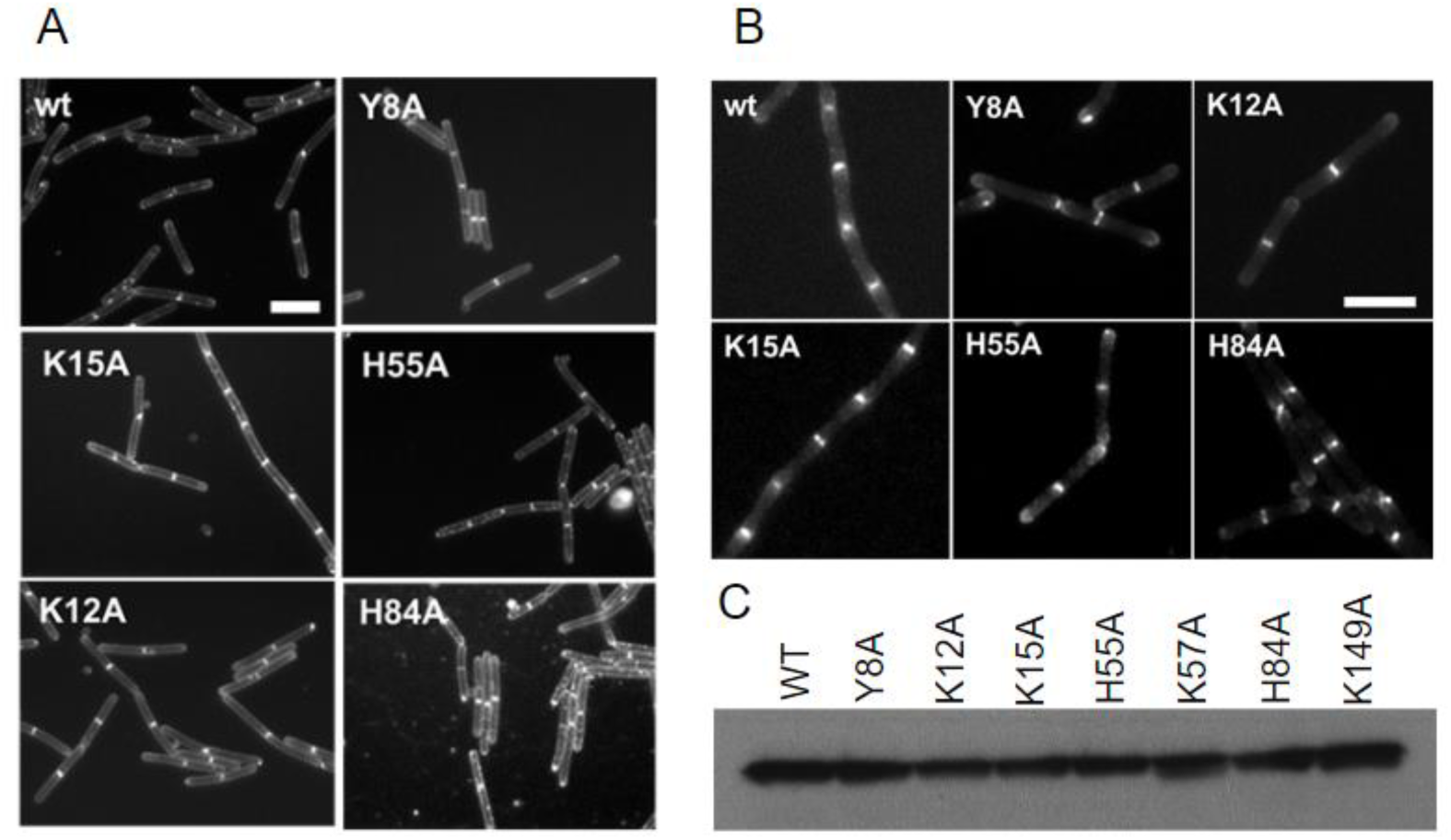
(A) Cell division phenotype of GFP-MinC^N^ mutants. All strains contain IPTG inducible alleles (Pspac) of GFP-MinC and were grown on LB pads with 0.2 mM of IPTG at 37°C for 3 hours. Cells were stained with FM 5-95. Scale bar 5 μm. (B) Subcellular localization of GFP-MinC^N^ mutants. Cells were induced with 0.2 mM IPTG, grown to exponential phase and immobilized on 1% agarose. Scale bar = 5 µm. (C) Western blot of whole cell extracts from each mutant revealed with anti-GFP antiserum. Blot contains 2 mutants (K57A and K149A) that are not relevant for this work and will be described elsewhere.

**Supplementary Figure 3.**
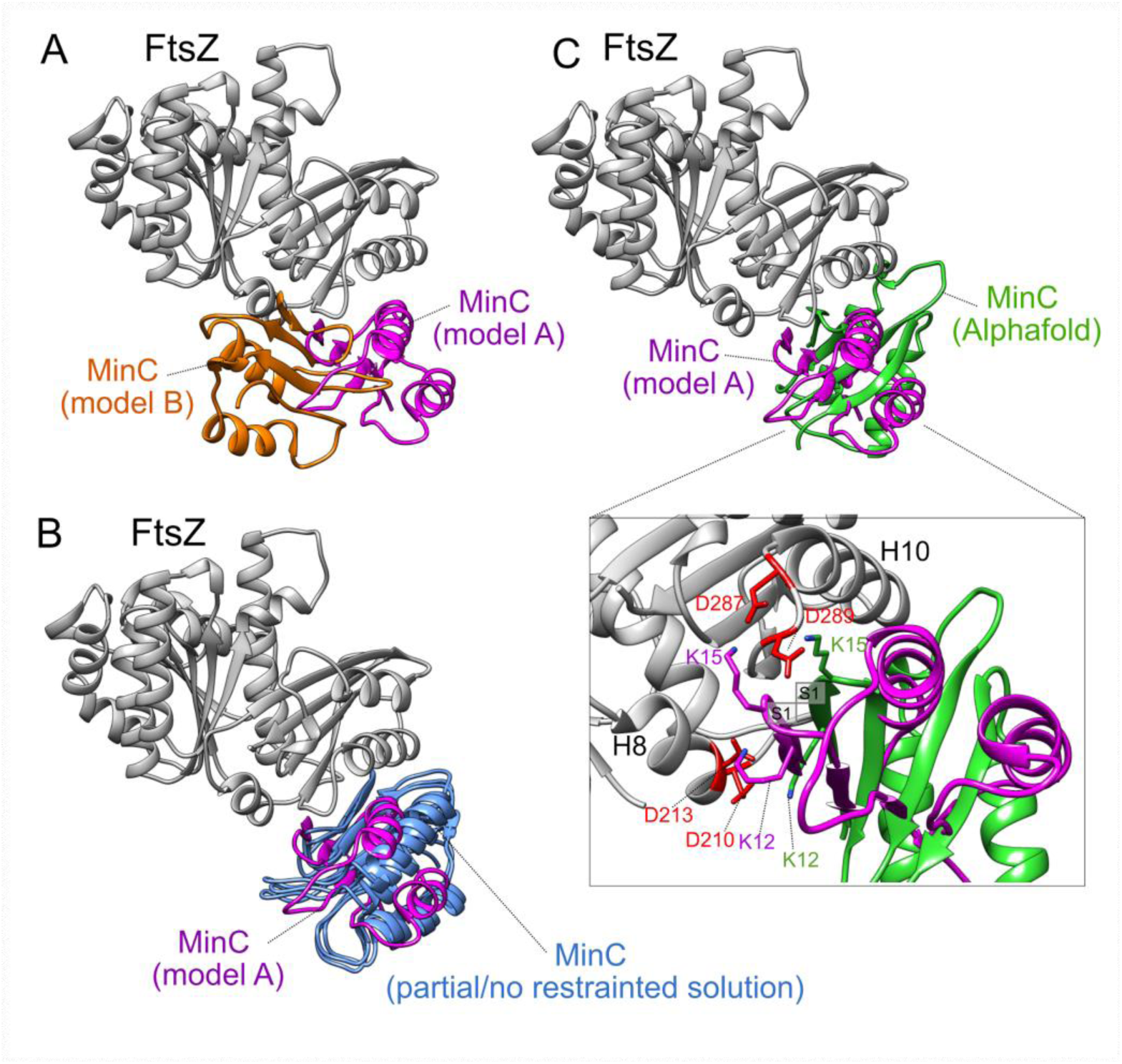
Comparison among 3D structural models of FtsZ:MinC complex. (A) Structural comparison between FtsZ:MinC 3D models (model A, in magenta, and model B, in orange) modeled by protein-protein docking with ClusPro server, using constraints based on experimental data. Models are aligned by FtsZ structure. (B) Structural comparison between model A (in magenta), generated in ClusPro server with constraints, and three other best ranked models (in blue), generated in ClusPro server with no constraints (Model C), with only MinC constraints (Model D), or with only FtsZ constraints (Model E). Since all these alternative models C, D and E, adopts a similar interaction position, we did not name them in the structure, for clarity. (C) Structural comparison between model A (in magenta), generated in ClusPro server with constraints, and the structural model generated by AlphaFold- Multimer (in green). In inset, a zoomed view of contacts between positively charged residues of MinC S1 and negatively charged residues of FtsZ. Important residues in this contact adopt similar positions in both proposed models.

**Supplementary Figure 4.**
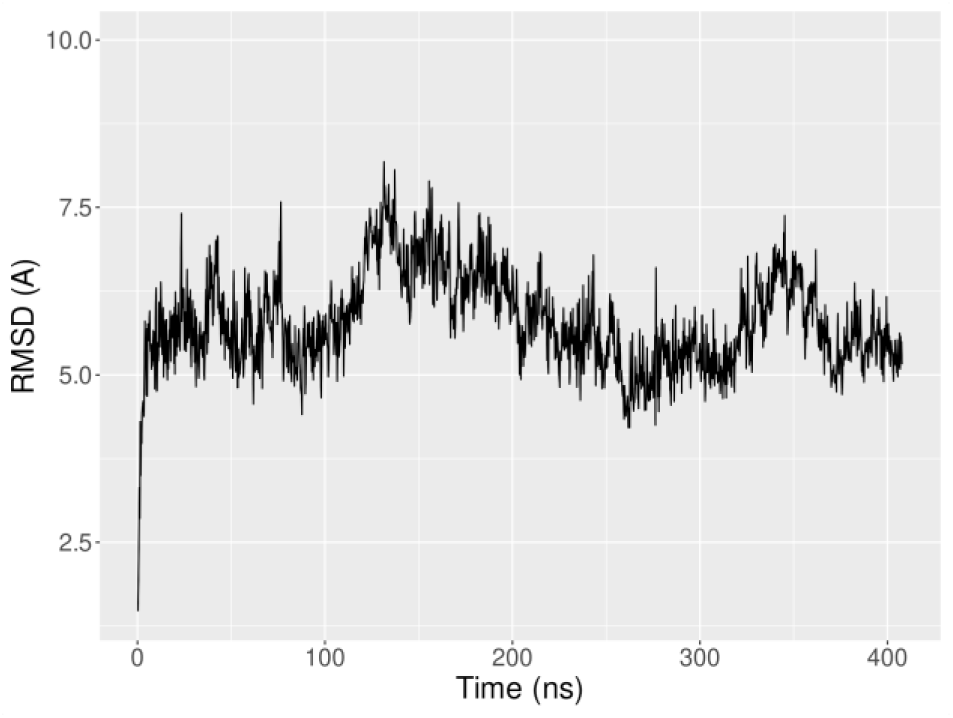
Root-mean-squared deviation (RMSD) of MinC during the 400 ns molecular dynamics simulation. The backbone MinC RMSD was calculated in VMD software and the system was aligned by FtsZ proteins using the first frame as reference.

**Supplemental Figure 5:**
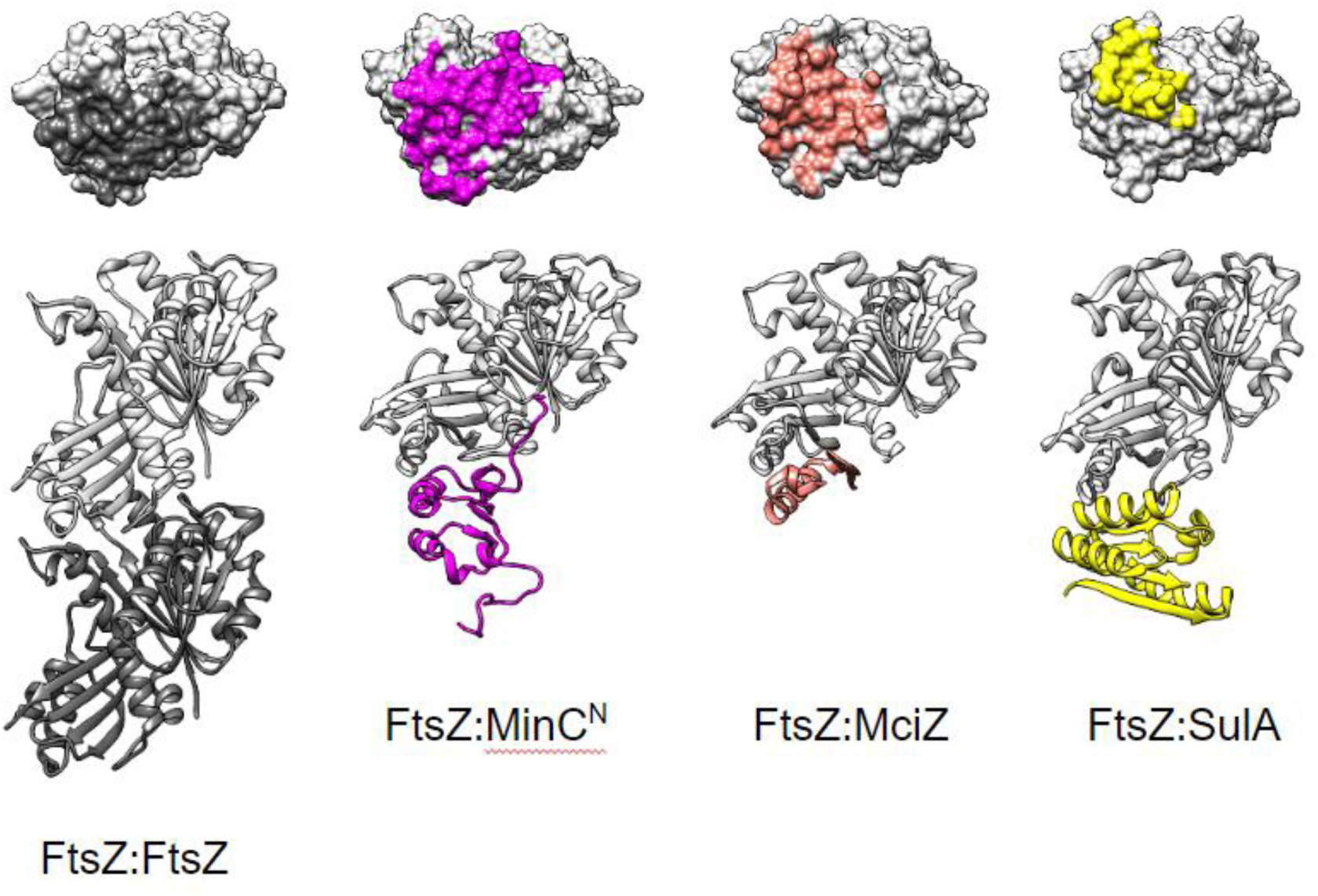
The C-terminal interface of FtsZ is a common target of negative modulators MinC, SulA and MciZ. From left to right: FtsZ dimer from the structure of a S. aureus FtsZ filament (5H5G); FtsZ-MinC^N^ complex predicted in this work (B. subtilis proteins); crystallographic structure of FtsZ-MciZ complex (4U39) (B. subtilis proteins); crystallographic structure of FtsZ-SulA complex (1OFU) (P. aeruginosa proteins). The top row shows contacts made by each partner with the FtsZ C-terminal (bottom, plus) polymerization face.

**Table S1.**
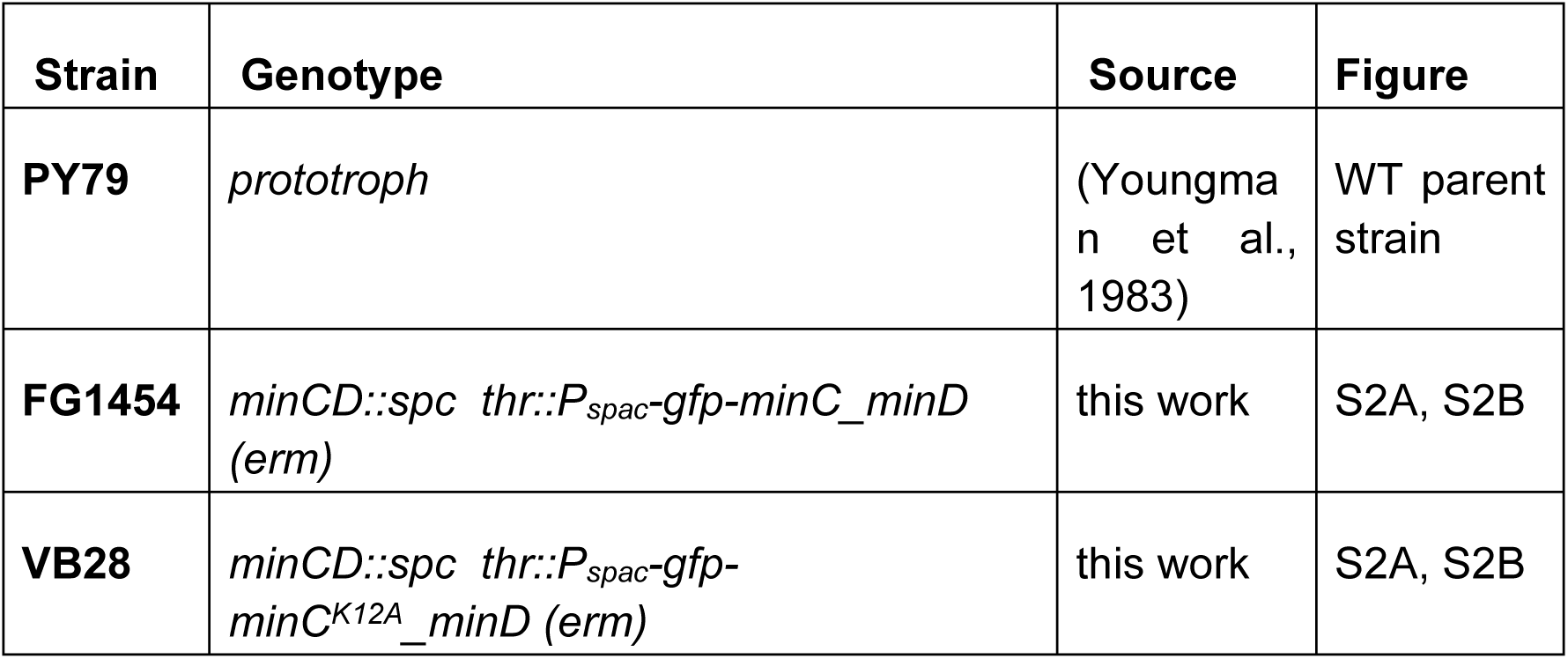
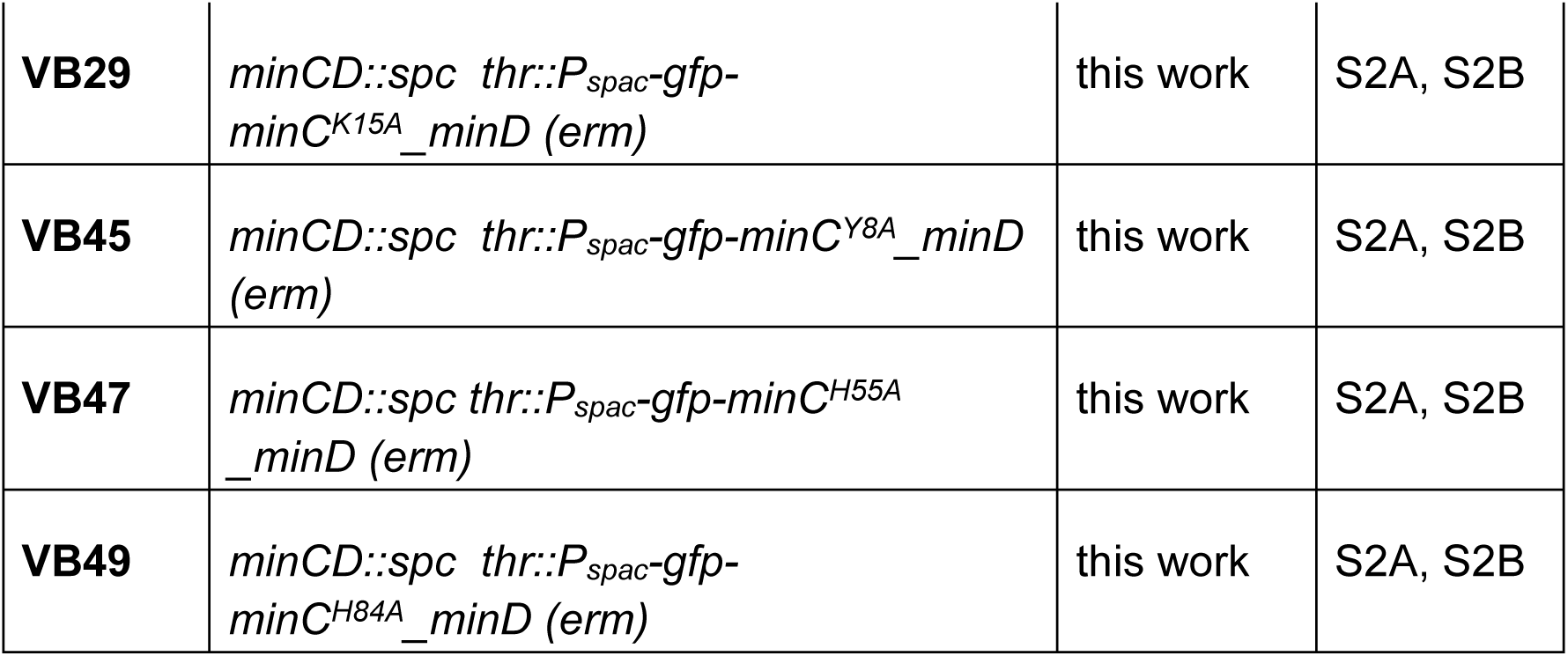
B. subtilis strains.

**Table S2.**
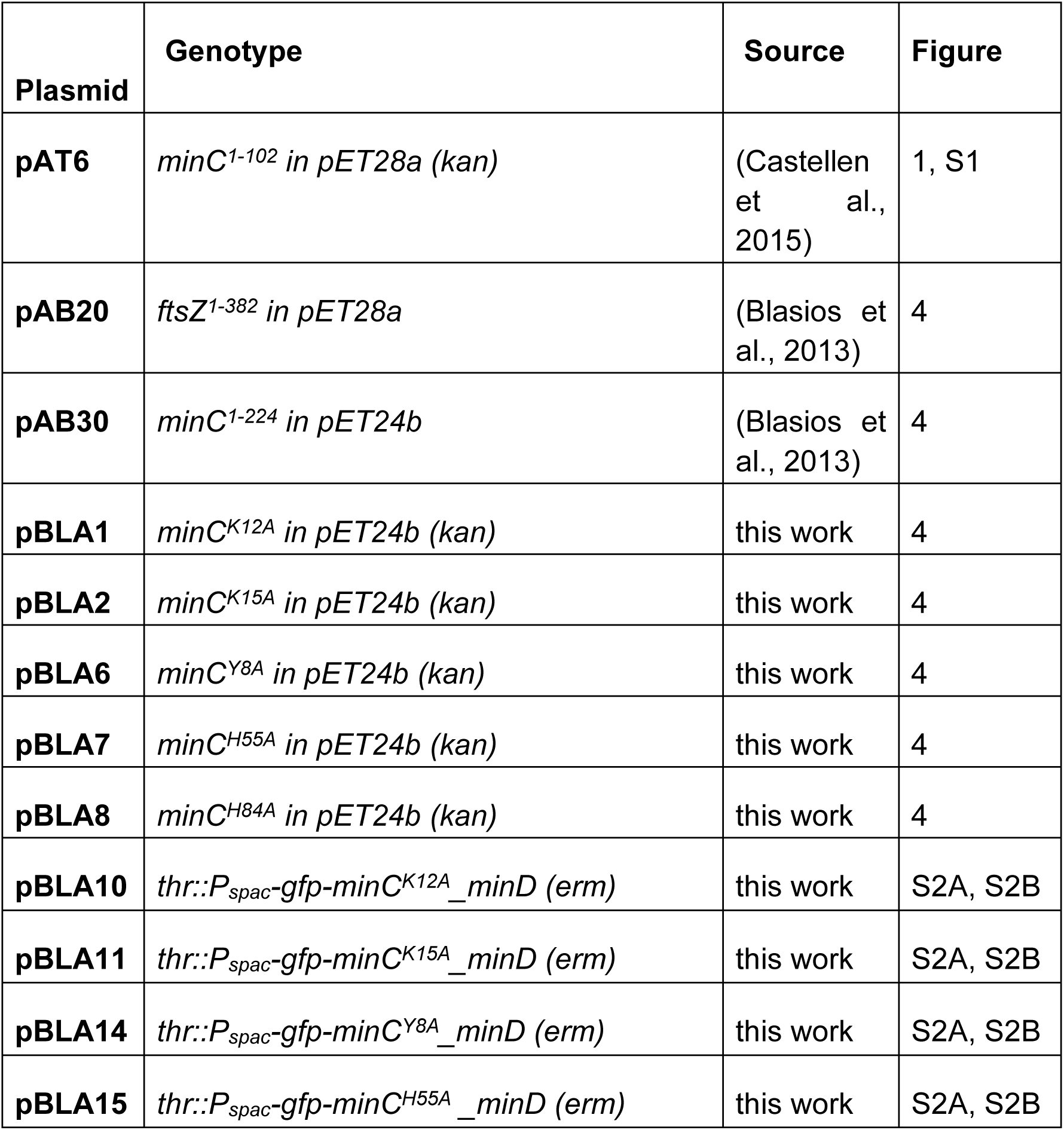
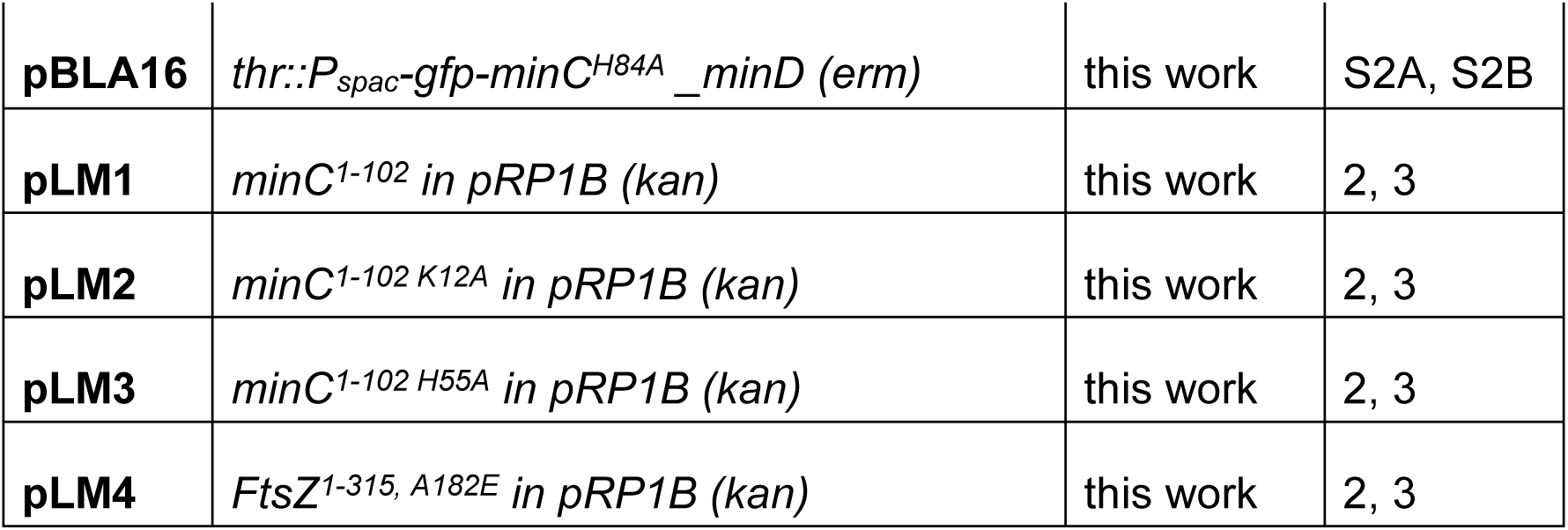
Plasmids.

**Table S3.** Frequency of hydrogen bond formation during MD simulation.

| MinC residues | FtsZ residues | HB frequency (%) |
| --- | --- | --- |
| LYS5-Main | ASN208-Side | 16.83% |
| LYS15-Side | ASP289-Side | 3.87% |
| LYS2-Side | ASP289-Side | 15.21% |
| TYR8-Side | ASP213-Side | 29.18% |
| LYS4-Side | LEU209-Main | 12.47% |
| LYS5-Side | GLY205-Main | 7.73% |
| SER2-Main | GLY37-Main | 17.21% |
| LYS12-Side | ASP213-Side | 26.31% |
| GLU42-Side | GLN288-Side | 8.73% |
| GLU42-Main | GLN288-Side | 3.99% |
| THR14-Side | GLN288-Main | 4.49% |
| LYS4-Main | ILE201-Main | 1.75% |
| LEU31-Main | GLN276-Side | 1.25% |

**Table S4.** Features of complex’s interfaces estimated using COCOMAPS (https://www.molnac.unisa.it/BioTools/cocomaps/). ^1^Note that our complex contains only MinC^N^ and FtsZ without its unstructured tail, whereas AlphaFold’s used the full length proteins.

| Parameter | NMR/Docking | AlphaFold |
| --- | --- | --- |
| Buried area upon complex formation (Å<sup>2</sup>) | 2353,6 | 2267,7 |
| Buried area upon complex formation (%) <sup>1</sup> | 11,41 | 5,77 |
| Interface area (Å<sup>2</sup>) | 1176,8 | 1133,85 |
| Interface area MOL1 (%) (MinC) <sup>1</sup> | 15,15 | 7,49 |
| Interface area MOL2 (%) (FtsZ) <sup>1</sup> | 9,15 | 4,69 |
| POLAR Buried area upon complex formation (Å<sup>2</sup>) | 1447,7 | 1536,2 |
| POLAR Interface (%) | 61,51 | 67,74 |
| POLAR Interface area (Å<sup>2</sup>) | 723,85 | 768,1 |
| NON POLAR Buried area upon complex formation (Å<sup>2</sup>) | 905,8 | 731,5 |
| NON POLAR Interface (%) | 38,49 | 32,26 |
| NON POLAR Interface area (Å<sup>2</sup>) | 452,9 | 365,75 |
| Residues at the interface_TOT (n) | 61 | 50 |
| Residues at the interface_Mol1 (MinC) | 25 | 22 |
| Residues at the interface_Mol2 (FtsZ) | 36 | 28 |
| Number of interacting residues Molecule1 (MinC) | 46 | 44 |
| Number of interacting residues Molecule2 (FtsZ) | 74 | 71 |
| Number of hydrophilic-hydrophobic interaction | 185 | 160 |
| Number of hydrophilic-hydrophilic interaction | 98 | 110 |
| Number of hydrophobic-hydrophobic interaction | 84 | 79 |

**Table S5.**
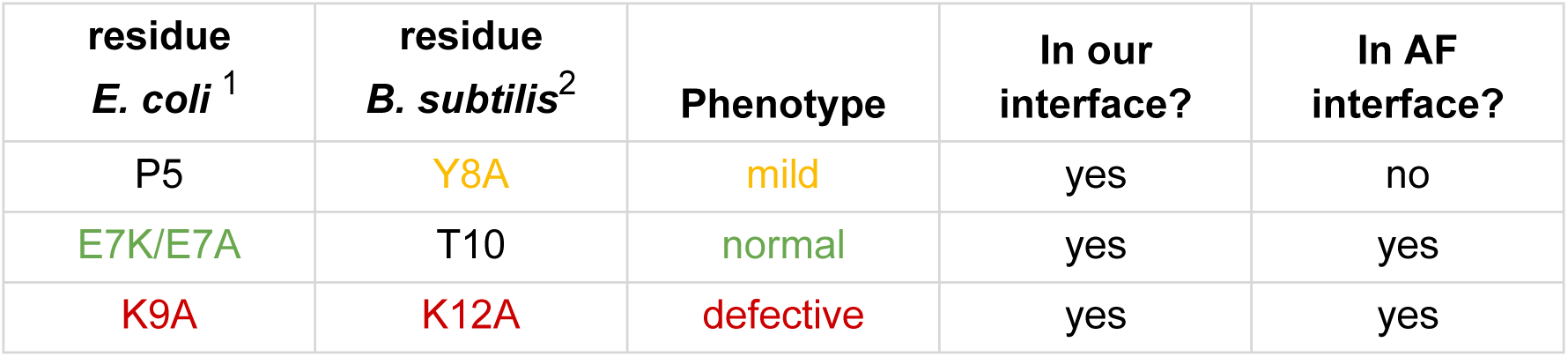
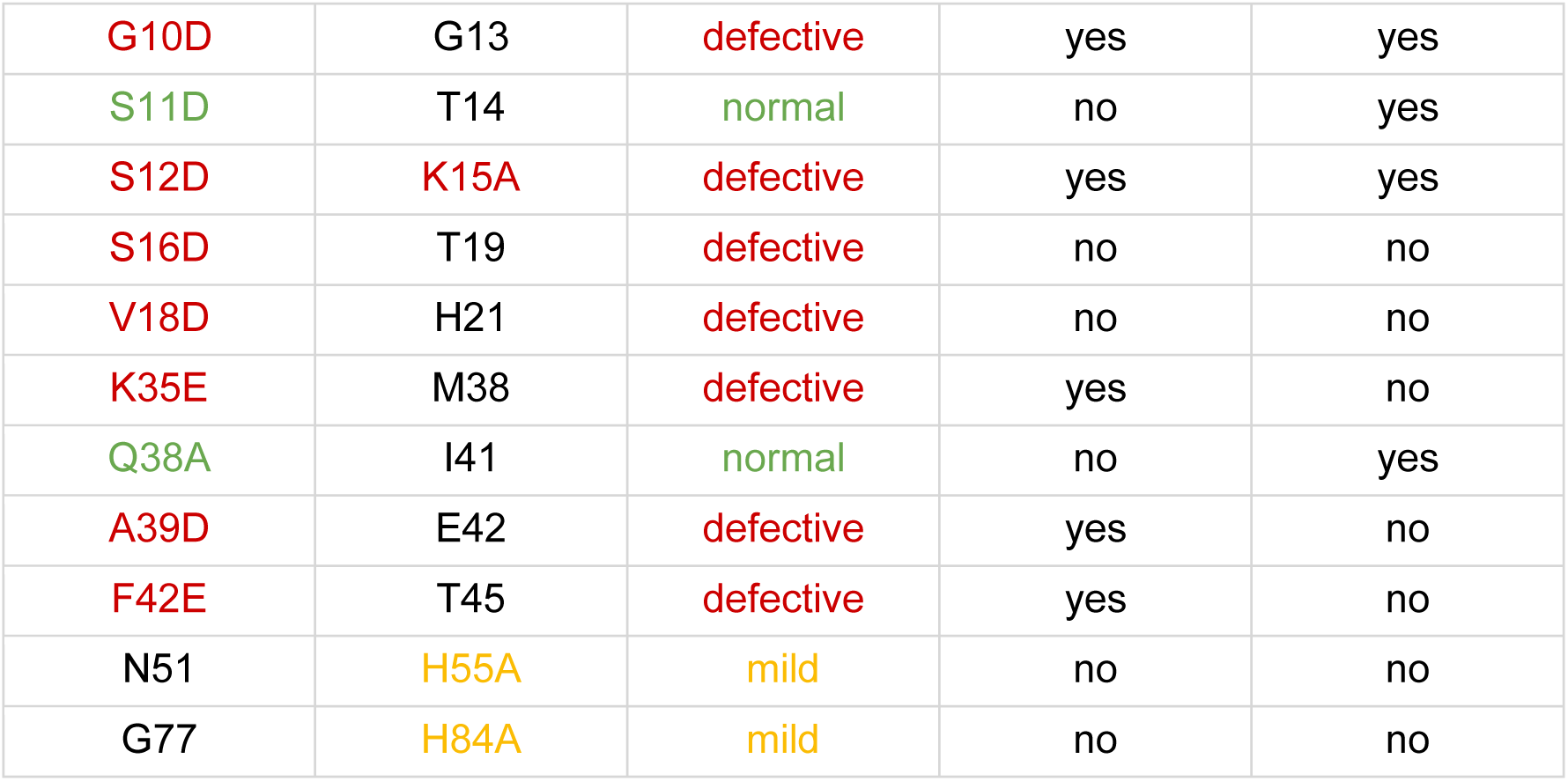
Comparison between MinC mutant data and residue presence in interface of predicted complexes. ^1^ Mutations in each species are colored according to phenotype. If a residue is not colored and has no substitution indicated it was not assayed in that species. ^2^ Residue correspondence as in Park et al. 2018.

| residue<br>*E. coli* <sup>1</sup> | residue<br>*B. subtilis* <sup>2</sup> | Phenotype | In our<br>interface? | In AF<br>interface? |
| --- | --- | --- | --- | --- |
| P5 | Y8A | mild | yes | no |
| E7K/E7A | T10 | normal | yes | yes |
| K9A | K12A | defective | yes | yes |
| G10D | G13 | defective | yes | yes |
| S11D | T14 | normal | no | yes |
| S12D | K15A | defective | yes | yes |
| S16D | T19 | defective | no | no |
| V18D | H21 | defective | no | no |
| K35E | M38 | defective | yes | no |
| Q38A | I41 | normal | no | yes |
| A39D | E42 | defective | yes | no |
| F42E | T45 | defective | yes | no |
| N51 | H55A | mild | no | no |
| G77 | H84A | mild | no | no |

## References

Adams, D.W., Wu, L.J., Errington, J., 2014. Cell cycle regulation by the bacterial nucleoid. Curr. Opin. Microbiol. 22, 94–101. 10.1016/j.mib.2014.09.020

An, J.Y., Kim, T.G., Park, K.R., Lee, J.-G., Youn, H.-S., Lee, Y., Kang, J.Y., Kang, G.B., Eom, S.H., 2013. Crystal structure of the N-terminal domain of MinC dimerized *via* domain swapping. J. Synchrotron Radiat. 20, 984– 988. 10.1107/S0909049513022760

Arumugam, S., Petrášek, Z., Schwille, P., 2014. MinCDE exploits the dynamic nature of FtsZ filaments for its spatial regulation. Proc. Natl. Acad. Sci. 111. 10.1073/pnas.1317764111

Bisson-Filho, A.W., Discola, K.F., Castellen, P., Blasios, V., Martins, A., Sforça, M.L., Garcia, W., Zeri, A.C.M., Erickson, H.P., Dessen, A., Gueiros-Filho, F.J., 2015. FtsZ filament capping by MciZ, a developmental regulator of bacterial division. Proc. Natl. Acad. Sci. U. S. A. 112, E2130–2138. 10.1073/pnas.1414242112

Blasios, V., Bisson-Filho, A.W., Castellen, P., Nogueira, M.L., Bettini, J., Portugal, R.V., Zeri, A.C., Gueiros-Filho, F.J., 2013. Genetic and biochemical characterization of the MinC-FtsZ interaction in Bacillus subtilis. PLoS One 8, e60690.

Bramkamp, M., Emmins, R., Weston, L., Donovan, C., Daniel, R.A., Errington, J., 2008. A novel component of the division site selection system of Bacillus subtilis and a new mode of action for the division inhibitor MinCD. Mol Microbiol 70, 1556–69.

Bramkamp, M., Van Baarle, S., 2009. Division site selection in rod-shaped bacteria. Curr. Opin. Microbiol. 12, 683–688. 10.1016/j.mib.2009.10.002

Castellen, P., Sforça, M.L., Gueiros-Filho, F.J., De Mattos Zeri, A.C., 2015. Backbone and side chain NMR assignments for the N-terminal domain of the cell division regulator MinC from Bacillus subtilis. Biomol. NMR Assign. 9, 1–5. 10.1007/s12104-013-9534-y

Chen, Y., Milam, S.L., Erickson, H.P., 2012. SulA Inhibits Assembly of FtsZ by a Simple Sequestration Mechanism. Biochemistry 51, 3100–3109. 10.1021/bi201669d

Clore, G.M., Venditti, V., 2013. Structure, dynamics and biophysics of the cytoplasmic protein–protein complexes of the bacterial phosphoenolpyruvate: sugar phosphotransferase system. Trends Biochem. Sci. 38, 515–530. 10.1016/j.tibs.2013.08.003

Corbin, L.C., Erickson, H.P., 2020. A Unified Model for Treadmilling and Nucleation of Single-Stranded FtsZ Protofilaments. Biophys. J. 119, 792– 805. 10.1016/j.bpj.2020.05.041

Cordell, S.C., Anderson, R.E., Lowe, J., 2001. Crystal structure of the bacterial cell division inhibitor MinC. EMBO J 20, 2454–61.

Cordell, S.C., Robinson, E.J., Lowe, J., 2003. Crystal structure of the SOS cell division inhibitor SulA and in complex with FtsZ. Proc Natl Acad Sci U A 100, 7889–94.

Corrales-Guerrero, L., Steinchen, W., Ramm, B., Mücksch, J., Rosum, J., Refes, Y., Heimerl, T., Bange, G., Schwille, P., Thanbichler, M., 2022. MipZ caps the plus-end of FtsZ polymers to promote their rapid disassembly. Proc. Natl. Acad. Sci. 119, e2208227119. 10.1073/pnas.2208227119

Dajkovic, A., Lan, G., Sun, S.X., Wirtz, D., Lutkenhaus, J., 2008a. MinC spatially controls bacterial cytokinesis by antagonizing the scaffolding function of FtsZ. Curr Biol 18, 235–44.

Dajkovic, A., Mukherjee, A., Lutkenhaus, J., 2008b. Investigation of regulation of FtsZ assembly by SulA and development of a model for FtsZ polymerization. J Bacteriol 190, 2513–26.

de Boer, P.A., Crossley, R.E., Rothfield, L.I., 1989. A division inhibitor and a topological specificity factor coded for by the minicell locus determine proper placement of the division septum in E. coli. Cell 56, 641–9.

de Oliveira, I.F., de Sousa Borges, A., Kooij, V., Bartosiak-Jentys, J., Luirink, J., Scheffers, D.J., 2010. Characterization of ftsZ mutations that render Bacillus subtilis resistant to MinC. PLoS One 5, e12048.

Delaglio, F., Grzesiek, S., Vuister, GeertenW., Zhu, G., Pfeifer, J., Bax, A., 1995. NMRPipe: A multidimensional spectral processing system based on UNIX pipes. J. Biomol. NMR 6. 10.1007/BF00197809

Du, S., Lutkenhaus, J., 2019. At the Heart of Bacterial Cytokinesis: The Z Ring. Trends Microbiol. 27, 781–791. 10.1016/j.tim.2019.04.011

Du, S., Pichoff, S., Kruse, K., Lutkenhaus, J., 2018. FtsZ filaments have the opposite kinetic polarity of microtubules. Proc. Natl. Acad. Sci. 115, 10768–10773. 10.1073/pnas.1811919115

Evans, R., O’Neill, M., Pritzel, A., Antropova, N., Senior, A., Green, T., Žídek, A., Bates, R., Blackwell, S., Yim, J., Ronneberger, O., Bodenstein, S., Zielinski, M., Bridgland, A., Potapenko, A., Cowie, A., Tunyasuvunakool, K., Jain, R., Clancy, E., Kohli, P., Jumper, J., Hassabis, D., 2021. Protein complex prediction with AlphaFold-Multimer (preprint). Bioinformatics. 10.1101/2021.10.04.463034

Fujita, J., Harada, R., Maeda, Y., Saito, Y., Mizohata, E., Inoue, T., Shigeta, Y., Matsumura, H., 2017. Identification of the key interactions in structural transition pathway of FtsZ from Staphylococcus aureus. J. Struct. Biol. 198, 65–73. 10.1016/j.jsb.2017.04.008

Galli, E., Poidevin, M., Le Bars, R., Desfontaines, J.-M., Muresan, L., Paly, E., Yamaichi, Y., Barre, F.-X., 2016. Cell division licensing in the multi-chromosomal Vibrio cholerae bacterium. Nat. Microbiol. 1, 16094. 10.1038/nmicrobiol.2016.94

Grant, B.J., Rodrigues, A.P.C., ElSawy, K.M., McCammon, J.A., Caves, L.S.D., 2006. Bio3d: an R package for the comparative analysis of protein structures. Bioinformatics 22, 2695–2696. 10.1093/bioinformatics/btl461

Gregory, J.A., Becker, E.C., Pogliano, K., 2008. Bacillus subtilis MinC destabilizes FtsZ-rings at new cell poles and contributes to the timing of cell division. Genes Dev 22, 3475–88.

Güntert, P., 2004. Automated NMR Structure Calculation With CYANA, in: Protein NMR Techniques. Humana Press, New Jersey, pp. 353–378. 10.1385/1-59259-809-9:353

Hammond, L.R., White, M.L., Eswara, P.J., 2019. ¡vIVA la DivIVA! J. Bacteriol. 201. 10.1128/JB.00245-19

Hernández-Rocamora, V.M., Alfonso, C., Margolin, W., Zorrilla, S., Rivas, G., 2015. Evidence That Bacteriophage λ Kil Peptide Inhibits Bacterial Cell Division by Disrupting FtsZ Protofilaments and Sequestering Protein Subunits. J. Biol. Chem. 290, 20325–20335. 10.1074/jbc.M115.653329

Hernández-Rocamora, V.M., García-Montañés, C., Reija, B., Monterroso, B., Margolin, W., Alfonso, C., Zorrilla, S., Rivas, G., 2013. MinC Protein Shortens FtsZ Protofilaments by Preferentially Interacting with GDP-bound Subunits. J. Biol. Chem. 288, 24625–24635. 10.1074/jbc.M113.483222

Hill, N.S., Buske, P.J., Shi, Y., Levin, P.A., 2013. A moonlighting enzyme links Escherichia coli cell size with central metabolism. PLoS Genet 9, e1003663.

Hu, Z., Lutkenhaus, J., 2000. Analysis of MinC reveals two independent domains involved in interaction with MinD and FtsZ. J Bacteriol 182, 3965–71.

Hu, Z., Mukherjee, A., Pichoff, S., Lutkenhaus, J., 1999. The MinC component of the division site selection system in Escherichia coli interacts with FtsZ to prevent polymerization. Proc Natl Acad Sci U A 96, 14819–24.

Johnson, B.A., Blevins, R.A., 1994. NMR View: A computer program for the visualization and analysis of NMR data. J. Biomol. NMR 4, 603–614. 10.1007/BF00404272

Kozakov, D., Hall, D.R., Xia, B., Porter, K.A., Padhorny, D., Yueh, C., Beglov, D., Vajda, S., 2017. The ClusPro web server for protein–protein docking. Nat. Protoc. 12, 255–278. 10.1038/nprot.2016.169

Krieger, E., Koraimann, G., Vriend, G., 2002. Increasing the precision of comparative models with YASARA NOVA—a self-parameterizing force field. Proteins Struct. Funct. Bioinforma. 47, 393–402. 10.1002/prot.10104

LaBreck, C.J., Conti, J., Viola, M.G., Camberg, J.L., 2019. MinC N- and C-Domain Interactions Modulate FtsZ Assembly, Division Site Selection, and MinD-Dependent Oscillation in *Escherichia coli*. J. Bacteriol. 201. 10.1128/JB.00374-18

Laskowski, Roman A., Rullmann, J. Antoon C., MacArthur, Malcolm W., Kaptein, R., Thornton, Janet M., 1996. AQUA and PROCHECK-NMR: Programs for checking the quality of protein structures solved by NMR. J. Biomol. NMR 8. 10.1007/BF00228148

Levin, P.A., Shim, J.J., Grossman, A.D., 1998. Effect of minCD on FtsZ ring position and polar septation in Bacillus subtilis. J Bacteriol 180, 6048–51.

Li, Y., Hsin, J., Zhao, L., Cheng, Y., Shang, W., Huang, K.C., Wang, H.-W., Ye, S., 2013. FtsZ Protofilaments Use a Hinge-Opening Mechanism for Constrictive Force Generation. Science 341, 392–395. 10.1126/science.1239248

Lorenzoni, A.S.G., Dantas, G.C., Bergsma, T., Ferreira, H., Scheffers, D.-J., 2017. Xanthomonas citri MinC Oscillates from Pole to Pole to Ensure Proper Cell Division and Shape. Front. Microbiol. 8, 1352. 10.3389/fmicb.2017.01352

Lutkenhaus, J., 2007. Assembly dynamics of the bacterial MinCDE system and spatial regulation of the Z ring. Annu Rev Biochem 76, 539–62.

MacCready, J.S., Schossau, J., Osteryoung, K.W., Ducat, D.C., 2017. Robust M in-system oscillation in the presence of internal photosynthetic membranes in cyanobacteria. Mol. Microbiol. 103, 483–503. 10.1111/mmi.13571

Marston, A.L., Thomaides, H.B., Edwards, D.H., Sharpe, M.E., Errington, J., 1998. Polar localization of the MinD protein of Bacillus subtilis and its role in selection of the mid-cell division site. Genes Dev 12, 3419–30.

McNally, F.J., Roll-Mecak, A., 2018. Microtubule-severing enzymes: From cellular functions to molecular mechanism. J. Cell Biol. 217, 4057–4069. 10.1083/jcb.201612104

McQuillen, R., Xiao, J., 2020. Insights into the Structure, Function, and Dynamics of the Bacterial Cytokinetic FtsZ-Ring. Annu. Rev. Biophys. 49, 309–341. 10.1146/annurev-biophys-121219-081703

Mullins, R.D., Heuser, J.A., Pollard, T.D., 1998. The interaction of Arp2/3 complex with actin: Nucleation, high affinity pointed end capping, and formation of branching networks of filaments. Proc. Natl. Acad. Sci. 95, 6181–6186. 10.1073/pnas.95.11.6181

Ono, S., 2007. Mechanism of Depolymerization and Severing of Actin Filaments and Its Significance in Cytoskeletal Dynamics, in: International Review of Cytology. Elsevier, pp. 1–82. 10.1016/S0074-7696(07)58001-0

Park, K.-T., Dajkovic, A., Wissel, M., Du, S., Lutkenhaus, J., 2018. MinC and FtsZ mutant analysis provides insight into MinC/MinD-mediated Z ring disassembly. J. Biol. Chem. 293, 5834–5846. 10.1074/jbc.M117.815894

Patrick, J.E., Kearns, D.B., 2008. MinJ (YvjD) is a topological determinant of cell division in Bacillus subtilis. Mol Microbiol 70, 1166–79.

Ramírez-Aportela, E., López-Blanco, J.R., Andreu, J.M., Chacón, P., 2014. Understanding Nucleotide-Regulated FtsZ Filament Dynamics and the Monomer Assembly Switch with Large-Scale Atomistic Simulations. Biophys. J. 107, 2164–2176. 10.1016/j.bpj.2014.09.033

Ramm, B., Heermann, T., Schwille, P., 2019. The E. coli MinCDE system in the regulation of protein patterns and gradients. Cell. Mol. Life Sci. 76, 4245– 4273. 10.1007/s00018-019-03218-x

Raskin, D.M., de Boer, P.A., 1999a. Rapid pole-to-pole oscillation of a protein required for directing division to the middle of Escherichia coli. Proc Natl Acad Sci U A 96, 4971–6.

Raskin, D.M., de Boer, P.A., 1999b. MinDE-dependent pole-to-pole oscillation of division inhibitor MinC in Escherichia coli. J Bacteriol 181, 6419–24.

Rowlett, V.W., Margolin, W., 2015. The Min system and other nucleoid-independent regulators of Z ring positioning. Front. Microbiol. 6. 10.3389/fmicb.2015.00478

Ruiz, F.M., Huecas, S., Santos-Aledo, A., Prim, E.A., Andreu, J.M., Fernández-Tornero, C., 2022. FtsZ filament structures in different nucleotide states reveal the mechanism of assembly dynamics. PLOS Biol. 20, e3001497. 10.1371/journal.pbio.3001497

Scheffers, D.J., 2008. The effect of MinC on FtsZ polymerization is pH dependent and can be counteracted by ZapA. FEBS Lett 582, 2601–8.

Schumacher, M.A., Ohashi, T., Corbin, L., Erickson, H.P., 2020. High-resolution crystal structures of *Escherichia coli* FtsZ bound to GDP and GTP. Acta Crystallogr. Sect. F Struct. Biol. Commun. 76, 94–102. 10.1107/S2053230X20001132

Shen, B., Lutkenhaus, J., 2010. Examination of the interaction between FtsZ and MinCN in E. coli suggests how MinC disrupts Z rings. Mol Microbiol 75, 1285–98.

Shen, B., Lutkenhaus, J., 2009. The conserved C-terminal tail of FtsZ is required for the septal localization and division inhibitory activity of MinC(C)/MinD. Mol Microbiol 72, 410–24.

Shiomi, D., Margolin, W., 2007. The C-terminal domain of MinC inhibits assembly of the Z ring in Escherichia coli. J Bacteriol 189, 236–43. Epub 2006 Nov 3.

Strahl, H., Hamoen, L.W., 2012. Finding the corners in a cell. Curr. Opin. Microbiol. 15, 731–736. 10.1016/j.mib.2012.10.006

Szeto, T.H., Rowland, S.L., King, G.F., 2001. The Dimerization Function of MinC Resides in a Structurally Autonomous C-Terminal Domain. J. Bacteriol. 183, 6684–6687. 10.1128/JB.183.22.6684-6687.2001

Terwilliger, T.C., Liebschner, D., Croll, T.I., Williams, C.J., McCoy, A.J., Poon, B.K., Afonine, P.V., Oeffner, R.D., Richardson, J.S., Read, R.J., Adams, P.D., 2024. AlphaFold predictions are valuable hypotheses and accelerate but do not replace experimental structure determination. Nat. Methods 21, 110–116. 10.1038/s41592-023-02087-4

Van Baarle, S., Bramkamp, M., 2010. The MinCDJ System in Bacillus subtilis Prevents Minicell Formation by Promoting Divisome Disassembly. PLoS ONE 5, e9850. 10.1371/journal.pone.0009850

Vranken, W.F., Boucher, W., Stevens, T.J., Fogh, R.H., Pajon, A., Llinas, M., Ulrich, E.L., Markley, J.L., Ionides, J., Laue, E.D., 2005. The CCPN data model for NMR spectroscopy: Development of a software pipeline. Proteins Struct. Funct. Bioinforma. 59, 687–696. 10.1002/prot.20449

Wagstaff, J.M., Tsim, M., Oliva, M.A., García-Sanchez, A., Kureisaite-Ciziene, D., Andreu, J.M., Löwe, J., 2017. A Polymerization-Associated Structural Switch in FtsZ That Enables Treadmilling of Model Filaments. mBio 8. 10.1128/mBio.00254-17

Wettmann, L., Kruse, K., 2018. The Min-protein oscillations in *Escherichia coli* : an example of self-organized cellular protein waves. Philos. Trans. R. Soc. B Biol. Sci. 373, 20170111. 10.1098/rstb.2017.0111

Yoshizawa, T., Fujita, J., Terakado, H., Ozawa, M., Kuroda, N., Tanaka, S., Uehara, R., Matsumura, H., 2020. Crystal structures of the cell-division protein FtsZ from *Klebsiella pneumoniae* and *Escherichia coli*. Acta Crystallogr. Sect. F Struct. Biol. Commun. 76, 86–93. 10.1107/S2053230X2000076X

Youngman, P.J., Perkins, J.B., Losick, R., 1983. Genetic transposition and insertional mutagenesis in Bacillus subtilis with Streptococcus faecalis transposon Tn917. Proc Natl Acad Sci U A 80, 2305–9.

Yu, Y., Zhou, J., Dempwolff, F., Baker, J.D., Kearns, D.B., Jacobson, S.C., 2020. The Min System Disassembles FtsZ Foci and Inhibits Polar Peptidoglycan Remodeling in Bacillus subtilis. mBio 11, e03197–19. 10.1128/mBio.03197-19

